# Inheritance of a Single Edited *CD46* Allele Is Associated with Reduced *Ex Vivo* Susceptibility to Bovine Viral Diarrhea Virus

**DOI:** 10.64898/2026.08.31.748238

**Authors:** Aspen M. Workman, Alexandria C. Krueger, Michael P. Heaton, Alexandria P. Snider, Kristen L. Kuhn, Tad S. Sonstegard, Brian L. Vander Ley

## Abstract

Bovine viral diarrhea virus (BVDV) remains an economically important pathogen of cattle despite widespread vaccination. A homozygous *CD46*-edited Gir heifer (Ginger) was previously shown to have significantly reduced susceptibility to BVDV. The edited allele contains an in-frame six amino acid substitution within the virus-binding domain of the BVDV entry receptor CD46, replacing residues G82QVLAL with A_82_LPTFS. Here, we investigated whether reduced BVDV susceptibility is maintained when the edited allele is inherited in the heterozygous state. Ginger was artificially inseminated with semen from an unedited Gir bull and produced a healthy heterozygous *CD46*-edited bull calf (Giraldo). Whole-genome sequencing confirmed the inheritance and structural integrity of Giraldo’s edited allele. Compared with Ginger, Giraldo exhibited similarly reduced *ex vivo* BVDV susceptibility across primary fibroblasts, lymphocytes, and monocytes, despite inheriting a wild-type *CD46* allele from the sire. Allele-specific *CD46* RNA expression analysis demonstrated expression of both the edited and wild-type *CD46* alleles. Thus, the reduced-susceptibility phenotype was not attributable to transcriptional silencing of the wild-type allele. Lentiviral complementation studies in *CD46*-knockout Madin-Darby bovine kidney (MDBK) cells further demonstrated that this wild-type *CD46* allele was competent to support BVDV infection when expressed independently. Together, these findings indicate that the *CD46* A_82_LPTFS allele can confer reduced BVDV susceptibility in the heterozygous state despite expression of a functional wild-type *CD46* allele. This result suggests the potential to more rapidly disseminate reduced BVDV susceptibility through conventional breeding using homozygous *CD46*-edited sires.

**Importance:** Bovine viral diarrhea virus (BVDV) continues to cause substantial economic losses to the cattle industry despite the availability of vaccines. We previously developed cattle homozygous for a precisely edited version of the BVDV entry receptor gene *CD46* that significantly reduces susceptibility to BVDV infection. Here, we show that this reduced-susceptibility phenotype is maintained when only one copy of the edited *CD46* gene is inherited, despite the presence of a functional unedited copy of the gene. This finding suggests that the edited allele can reduce BVDV susceptibility in heterozygous animals, making it feasible to disseminate this trait through conventional breeding. More broadly, this study demonstrates the potential for precise editing of host genes exploited by viruses to generate heritable disease resistance, offering a practical strategy to improve livestock health and resilience.

## Introduction

Bovine viral diarrhea virus (BVDV) is a globally important pathogen and remains one of the most economically consequential viral diseases affecting the cattle industry (1–3). These economic losses result from a combination of respiratory and enteric disease, reproductive failure, and immunosuppression. Reproductive losses are particularly consequential, as the virus can cause fetal death, abortion, and the birth of persistently infected (PI) calves, which become a lifelong source of infection for the herd (4). The immunosuppressive effects also make cattle more vulnerable to other common diseases, including bovine respiratory disease complex (BRDC), thus compounding the economic impact. The persistence of these widespread impacts is striking given that vaccines against BVDV have been available for over 60 years (5, 6). This reality underscores the need for complementary or alternative prevention and control strategies.

Gene editing offers a complementary, host-directed genetic strategy for BVDV prevention. BVDV enters bovine cells through the host-encoded receptor CD46 (7). Our recent study demonstrated that a precise genome edit in the *CD46* gene led to significantly reduced BVDV susceptibility in an unvaccinated Gir heifer (“Ginger”) that was homozygous for the edited allele (8). This edit was introduced with CRISPR/Cas9-mediated homology-directed repair (HDR) to generate a 19-nucleotide in-frame substitution in exon 2 of the *CD46* gene. This edit replaced the highly conserved CD46 motif G_82_QVLAL, previously identified as essential for BVDV binding, with the sequence A_82_LPTFS (9). Across MDBK cells, Gir fetal cell lines, and Ginger, we demonstrated that homozygosity for this edited *CD46* allele consistently impaired viral entry resulting in protection from BVDV infection in fetal tissues and during an *in vivo* challenge study (8). More recently, the identical A_82_LPTFS edit conferred reduced *ex vivo* BVDV susceptibility in primary cells derived from cloned American Angus calves, demonstrating that this strategy is effective across genetically distinct cattle populations (10).

Beyond its role in BVDV entry, CD46 has a well-established role as a complement regulatory molecule that protects host cells from autologous complement-mediated damage (11, 12). CD46 has also been suggested to play a role in mammalian fertilization (13, 14), although the biological significance of this function remains less clearly defined. Therefore, evaluating potential unintended consequences of the *CD46* edit remains essential, including effects on established CD46 functions and reproductive fitness. A recent study by Snider et al. (2025) demonstrated that oocytes from Ginger showed normal fertilization rates during *in vitro* fertilization with semen from an unedited cross-bred beef bull (15). Furthermore, Ginger was successfully artificially inseminated with semen from an unedited purebred Gir bull. Pregnancy was confirmed at day 35 of gestation, as previously reported (15), and the pregnancy subsequently progressed to full term, resulting in the birth of a healthy bull calf (“Giraldo”). These findings suggest that the *CD46* edit does not impair fertility or reproductive function in this homozygous *CD46*-edited female.

Although homozygous *CD46* A_82_LPTFS editing confers significantly reduced susceptibility to BVDV, it remained unknown whether a single edited allele would be sufficient to confer this phenotype, as expression of the unedited allele could potentially provide functional CD46 receptor activity. To address this question, BVDV susceptibility was investigated in primary cells derived from Giraldo, Ginger’s heterozygous offspring. Inheritance of a single edited *CD46* allele was associated with *ex vivo* reductions in BVDV susceptibility comparable to those observed in the homozygous edited heifer, Ginger, despite inheritance of a wild-type *CD46* allele from the sire. These findings provide a basis for evaluating the potential of this edit to facilitate dissemination of reduced BVDV susceptibility through conventional cattle breeding.

## Results

### Birth and Genotypic Validation of the *CD46* Heterozygous Calf

Ginger, a homozygous *CD46*-edited Gir heifer, was artificially inseminated with pooled semen from two wild-type Gir sires. This breeding resulted in the birth of a healthy bull calf, Giraldo, on December 7, 2024. Whole-genome sequencing identified Zacarias as the sire based on exclusion analysis using genotypes at 115 previously validated parentage SNPs **(Table S1)**. The sequence data further confirmed Giraldo’s expected heterozygous *CD46* genotype and verified the integrity of both the wild-type (wt, G_82_QVLAL) and edited (e, A_82_LPTFS) *CD46* alleles **(Fig. 1)**.

**Figure 1.**
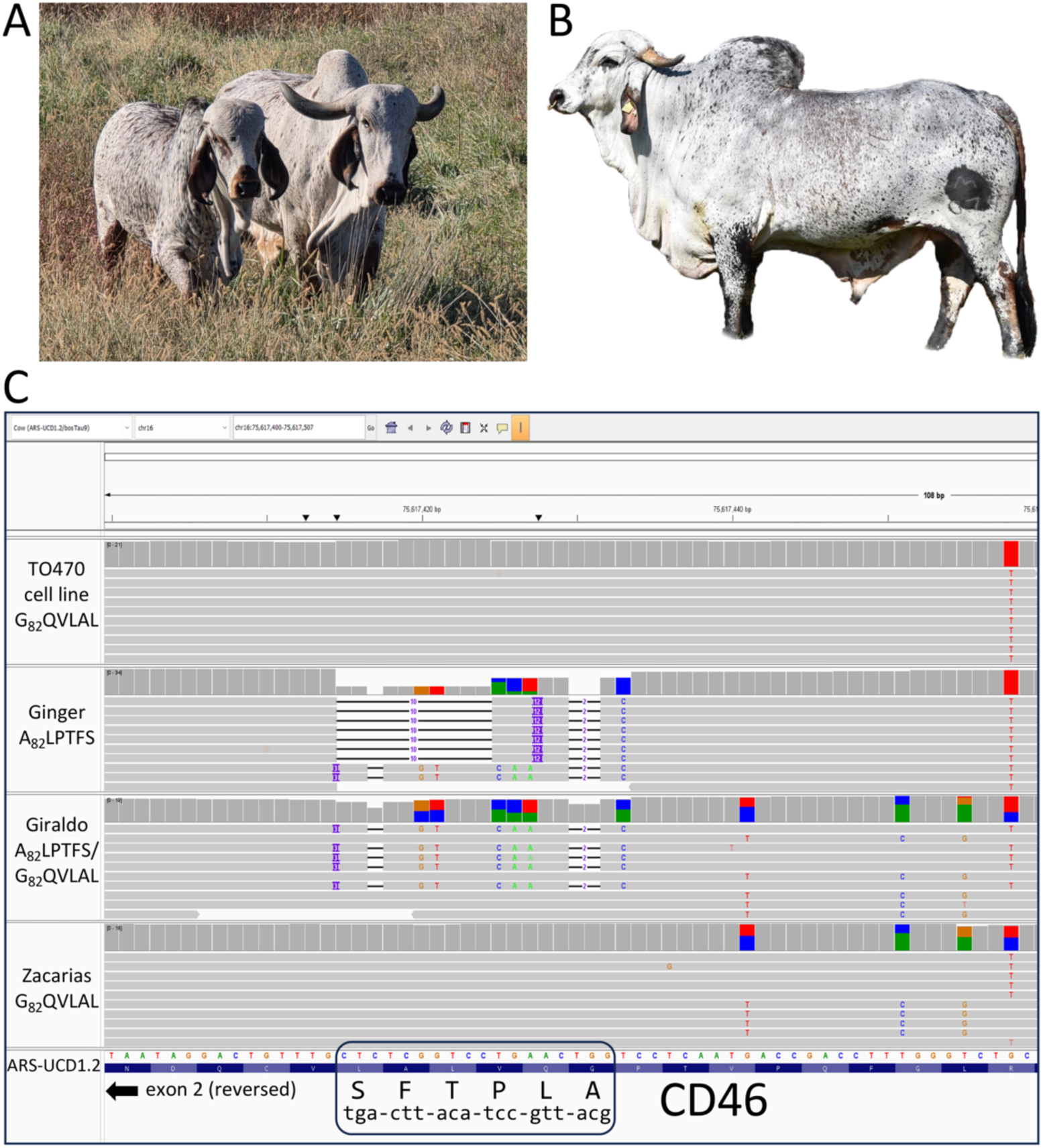
WGS confirmation of *CD46* genotypes from Giraldo’s pedigree. **A.** The edited dam (Ginger, right) and her bull calf (Giraldo, 10 months old). **B.** The sire (Zacarias). **C.** Screenshot showing the WGS paired-end data for each sample aligned to the relevant *CD46* exon 2 region harboring the A_82_LPTFS edit. Samples from top to bottom: the unedited TO470 cell line (homozygous G_82_QVLAL) used to produce Ginger; Ginger (homozygous edited A_82_LPTFS); Giraldo (heterozygous edited A_82_LPTFS); and Zacarias (homozygous G_82_QVLAL). The session file can be loaded directly into IGV for further viewing at the following URL: https://s3.us-west-2.amazonaws.com/usmarc.heaton.public/WGS/CellLines/ARS1.2/sessions/ARS1.2_CD46_TO470GingerGerryZac4Tracks.xml.

### CD46 Expression and BVDV Susceptibility in Primary Cells from *CD46*-Edited Cattle

CD46 protein expression and localization were first assessed in primary skin fibroblasts to confirm CD46 abundance and cellular distribution. Primary fibroblasts were analyzed from four animals representing three *CD46* genotypes: the parental unedited Gir fibroblast line used to clone Ginger (TO470, wild-type/wild-type [wt/wt]), a homozygous *CD46*-edited Gir dam (Ginger, edited/edited [e/e]), a heterozygous *CD46*-edited Gir offspring (Giraldo, edited/wild-type [e/wt]), and an unedited Holstein control (Mary Ann, wt/wt). There was no statistically significant difference in CD46 protein expression among TO470, Ginger, and Giraldo cells (**Fig. 2A**). Fibroblasts from Mary Ann showed a modest but statistically significant 1.3-fold increase in CD46 expression compared with TO470 cells (*p* < 0.05). Immunofluorescence analysis further demonstrated that CD46 localization was unchanged by the edit (**Fig. 2B**). Thus, the *CD46* edit, whether present in one or two copies, did not significantly alter CD46 protein expression or localization in primary skin fibroblasts.

**Figure 2.**
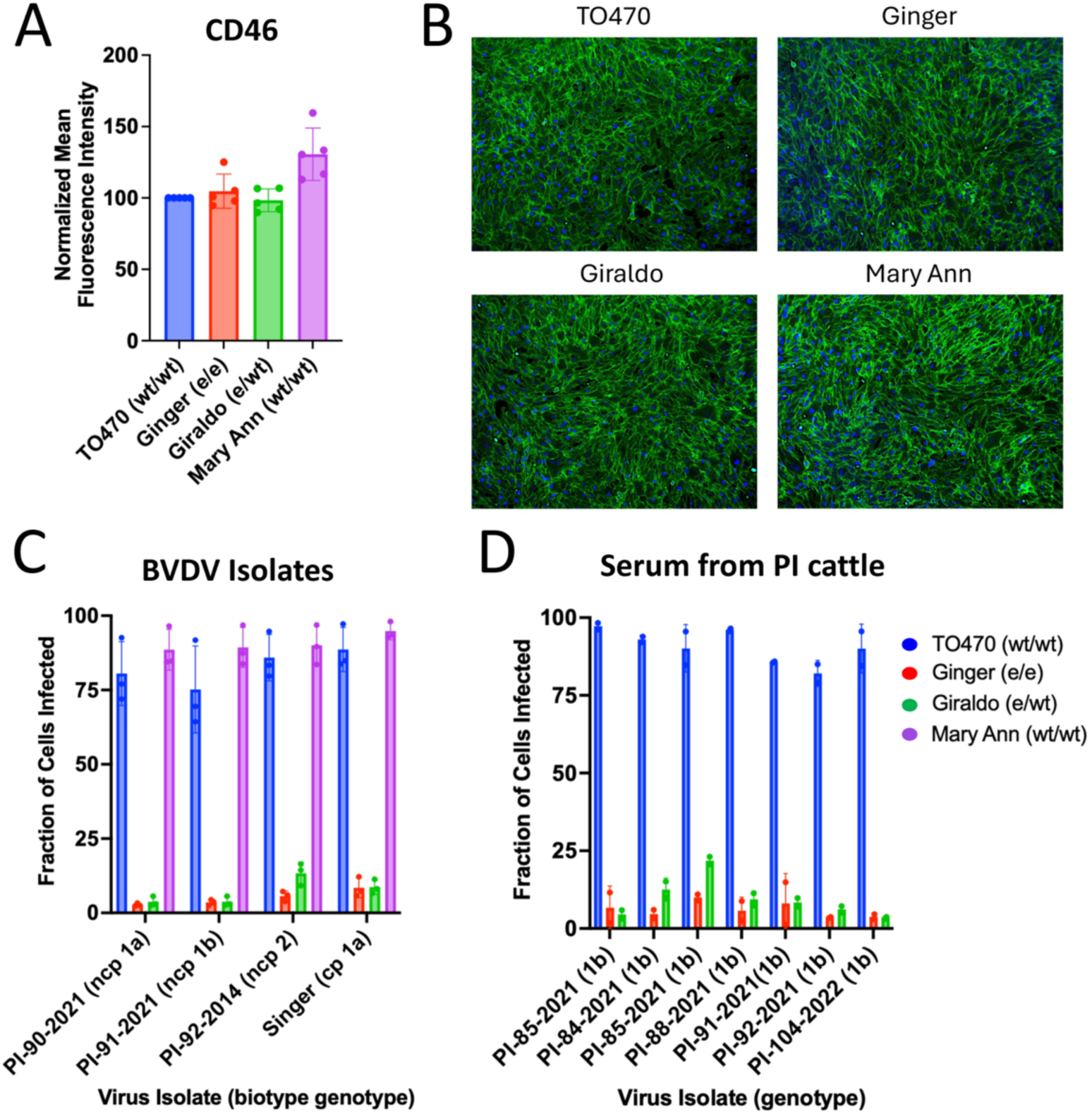
CD46 expression and *ex vivo* BVDV susceptibility in primary skin fibroblasts. **A.** Flow cytometric quantification of CD46 protein expression in uninfected cells with levels normalized to TO470. Results represent the mean ± standard deviation of *n* = 5 independent experiments. **B.** Immunofluorescence staining of CD46 (FITC) and nuclei were stained with DAPI (10x magnification). **C.** Cells were infected with cytopathic (cp) or non-cytopathic (ncp) BVDV isolates and infection efficiency was determined at 20 hpi by flow cytometry using a monoclonal anti-BVDV E2 antibody. Results represent the mean ± standard deviation of *n* = 3 independent experiments. **D.** Cells were inoculated with serum from BVDV PI cattle, and infection efficiency was quantified at 72 hpi by flow cytometry using an anti-BVDV E2 antibody. Results represent the mean ± standard deviation of *n* = 2 independent experiments. Abbreviations: wt, wild-type (unedited) allele; e, edited allele.

BVDV susceptibility was next assessed in these fibroblasts. Cells were challenged with isolates representing both major BVDV genotypes (BVDV-1 and BVDV-2) and biotypes (cytopathic and non-cytopathic). BVDV infection was significantly reduced in cells from both Ginger and Giraldo compared to wild-type controls (*p* < 0.05; **Fig. 2C**). The magnitude of this reduction was broadly comparable between Ginger and Giraldo, with BVDV susceptibility reduced by an average of 94% in Ginger cells (range: 91–97%) and 91% in Giraldo cells (range: 85–95%). Similarly, inoculation of fibroblasts with serum from persistently infected cattle demonstrated an average reduction in susceptibility of 93% for Ginger cells (range: 89–96%) and 90% for Giraldo cells (range: 76–96%) (**Fig. 2D**). A significant difference between Ginger and Giraldo was observed for a single serum sample (PI-85-2021; *p*=0.02), with greater susceptibility in Giraldo cells; however, this difference was small relative to the overall magnitude of the reduction observed in both edited animals compared with the wild-type control. Together, these findings show that primary fibroblasts from the heterozygous calf Giraldo exhibit a reduction in BVDV susceptibility comparable to that observed in the homozygous edited animal, despite the presence of a wild-type *CD46* allele.

BVDV susceptibility was subsequently assessed in primary lymphocytes and monocytes, which are important cellular targets for BVDV infection and dissemination. A sex-and age-matched crossbred taurine beef bull calf, “Bueno”, was used as a source of unedited control cells. Lymphocytes from Ginger and Giraldo showed significantly reduced susceptibility to BVDV compared to Bueno (*p* < 0.05), with the greatest reduction in susceptibility observed against BVDV-1 genotypes (**Fig. 3A**). On average, Ginger’s lymphocytes showed a 95% reduction in susceptibility to BVDV-1 (range: 90–99%) and a 63% reduction to BVDV-2 (range: 62–66%). Similarly, Giraldo’s lymphocytes demonstrated an average reduction of 93% to BVDV-1 (range: 85–98%) and 78% to BVDV-2 (range: 77–79%). Although Giraldo lymphocytes showed significantly lower susceptibility than Ginger lymphocytes for two isolates (PI-92-2014 and PI-65-2014; *p* < 0.05), overall lymphocyte susceptibility was highly comparable between Ginger and Giraldo across the 11 BVDV isolates tested.

**Figure 3.**
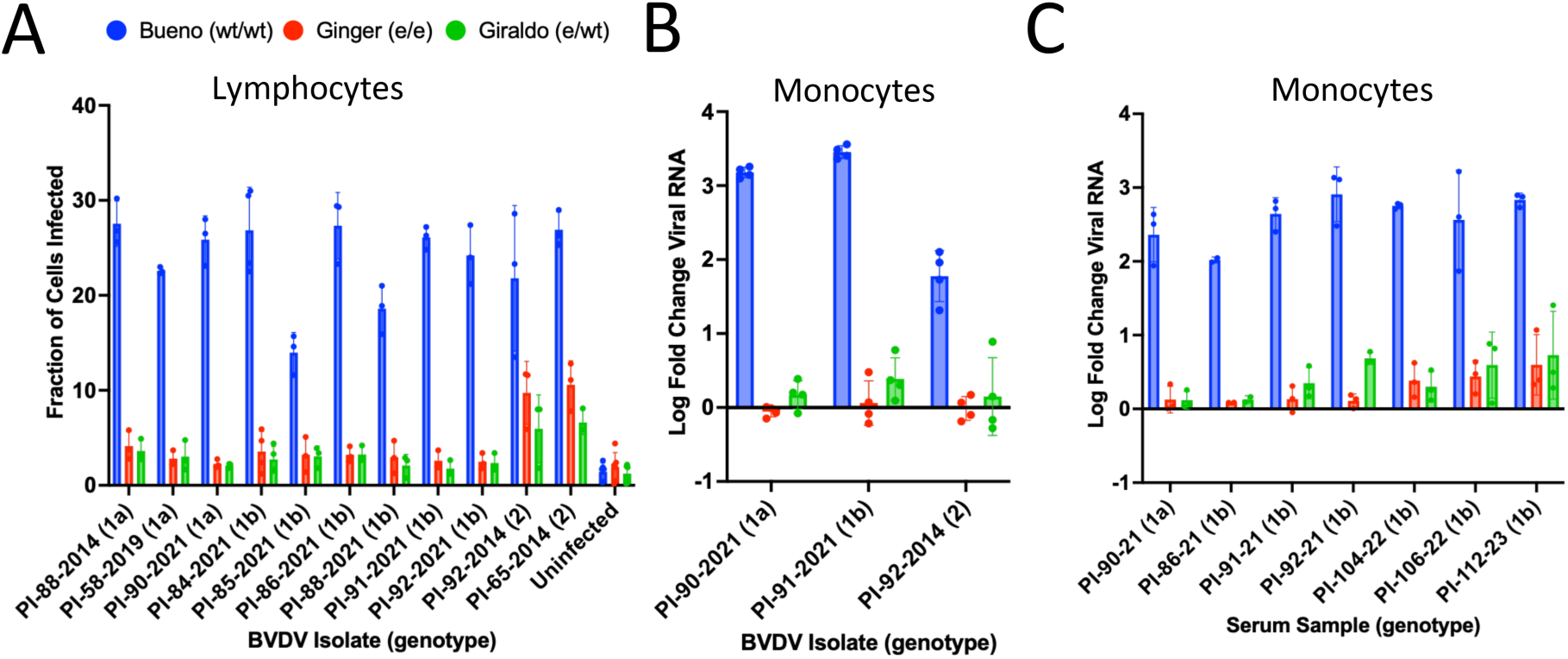
Ex vivo BVDV susceptibility in white blood cells from *CD46*-edited cattle. **A.** Flow cytometric quantification of BVDV-infected cells at 32 hpi. Results represent the mean ± SD of *n* = 3 independent experiments. **B.** Monocytes were infected with BVDV at a low MOI. At 72 hpi, viral RNA was detected by RT-qPCR and fold change in viral RNA relative to the input sample (0 hpi) was calculated. Results represent the mean ± SD of *n* = 3 independent experiments. **C.** Monocytes were inoculated with serum from BVDV-PI cattle and infection efficiency was determined at 48 hpi by RT-qPCR and fold change in viral RNA relative to the input sample (0 hpi) was calculated. Results represent the mean ± SD of at least *n* = 3 independent experiments. Abbreviations: wt, wild-type (unedited) *CD46* allele; e, edited *CD46* allele.

Monocytes from the *CD46*-edited cattle, Ginger and Giraldo, had a significant and comparable reduction in susceptibility to BVDV as quantified by RT-qPCR. Across the three BVDV isolates tested, viral RNA replication was reduced by an average of 95% compared to the wild-type control (**Fig. 3B**). No significant difference in susceptibility was observed between Ginger and Giraldo for any of the three BVDV isolates tested. When monocytes were inoculated with serum from persistently infected cattle, both Ginger and Giraldo cells showed significantly reduced BVDV susceptibility compared to the wild-type control across all seven serum samples tested (**Fig. 3C**). Across these samples, Ginger monocytes exhibited an average 90% reduction in susceptibility (range: 79–96%), while Giraldo monocytes showed an average 85% reduction (range: 74–95%). A significant difference between Ginger and Giraldo was observed for a single serum sample (PI-92-2021), with greater susceptibility in Giraldo monocytes. Together, these results demonstrate that peripheral blood cells from animals with one or two edited *CD46* alleles exhibit comparable reductions in BVDV susceptibility across diverse viral challenges.

### Allele-Specific Expression of *CD46* in the Heterozygous Calf

Given that Giraldo exhibited reduced BVDV susceptibility comparable to that of his homozygous *CD46*-edited dam, we next investigated whether expression of the wild-type allele was altered in Giraldo and contributed to this phenotype. Allele-specific *CD46* expression was assessed using both long-read, full-length RNA transcript sequencing and short-read RNA sequencing of RNA isolated from Giraldo’s whole blood and primary skin fibroblasts. Long-read sequencing generated 15.9 million circular consensus sequences (CCS) from primary skin fibroblasts and 14.3 million CCS from whole blood, while short-read sequencing generated 40.3 million and 73.7 million paired-end reads, respectively.

Long-read transcript analysis revealed that the edited *CD46* allele was expressed at a significantly higher level than the expected 1:1 allelic ratio in both skin fibroblasts (53.4% edited vs. 46.6% wt; *p* < 0.05) and whole blood (59.3% vs. 40.7%; *p* < 0.001) (**Table 1**). Short-read RNA sequencing independently showed the same pattern, with a bias toward expression of the edited allele in both skin fibroblasts (58.5% vs. 41.5%; *p* < 0.001) and whole blood (54.3% vs. 45.7%; *p* < 0.05) (**Table 1**). The concordance between the two sequencing approaches provides independent evidence that both the edited and wild-type *CD46* alleles are expressed in Giraldo’s cells. Thus, the reduced BVDV susceptibility phenotype cannot be attributed simply to complete loss of wild-type *CD46* allele expression, although the modest and reproducible bias toward expression of the edited allele may contribute to the observed phenotype.

**Table 1.** Quantification of edited and wild-type *CD46* allele expression in Giraldo’s primary cells. Asterisks denote a statistically significant deviation from an expected 1:1 allelic ratio, determined by Pearson’s chi-square goodness-of-fit test (*p < 0.05; **p < 0.001).

| Sample | Allele Variant | Long-Read mRNA |  | Short-read mRNA |  |
| --- | --- | --- | --- | --- | --- |
|  |  | Reads (CD46) | Percentage (%) | Allele-specific read count | Percentage (%) |
| Skin Cells | Edited (A<sub>82</sub>LPTFS) | 535 | 53.4* | 231 | 58.5** |
|  | Wild-type (G<sub>82</sub>QVLAL) | 467 | 46.6 | 164 | 41.5 |
|  | **Total** | 1002 | 100 | 395 | 100 |
| Blood Cells | Edited (A<sub>82</sub>LPTFS) | 198 | 59.3** | 299 | 54.3* |
|  | Wild-type (G<sub>82</sub>QVLAL) | 136 | 40.7 | 251 | 45.7 |
|  | **Total** | 334 | 100 | 550 | 100 |

### Functional Characterization of *CD46* Alleles Using a Lentiviral Complementation Model

Having established that both edited and wild-type *CD46* alleles are expressed in the heterozygous bull calf, we next determined whether the paternally inherited wild-type Zacarias *CD46* allele encodes a functional receptor capable of supporting BVDV infection. Comparison of the predicted Zacarias-derived CD46 protein sequence with that of native MDBK CD46 identified 13 amino acid polymorphisms, corresponding to 96.5% amino acid sequence identity (**Table 2**). Two substitutions were of particular interest because V69L occurs within, and Y70H immediately adjacent to, the E_66_QIV_69_ motif located within the first complement control protein repeat (CCP1) domain, which contributes to the BVDV binding interface (9). Previous work has shown that naturally occurring amino acid variation within the CCP1 domain can reduce CD46-mediated BVDV entry, indicating that sequence differences in this region may influence receptor function (16).

**Table 2.** Non-synonymous single nucleotide polymorphisms (SNPs) distinguishing the Zacarias *CD46* allele and the Ginger *CD46* allele from native MDBK *CD46*. The bovine CD46 protein has four N-terminal complement control protein (CCP) domains (CCP1-4), a transmembrane domain, and a C-terminal cytoplasmic tail (CYT) involved in intracellular signaling. The BVDV binding platform is located within the CCP1 domain and is composed of amino acids E_66_QIV_69_ and G_82_QVLAL_87_ (9). Highlighted are the non-synonymous SNPs that differentiate the Zacarias allele from the native MDBK CD46 allele. Amino acid positions are relative to the *Bos indicus* Genbank accession XP_070624793.1 and *Bos taurus* accession NP_001229491.2.

| Animal | Protein domain (amino acid positions) |  |  |  |  |  |  |  |  |  |  |  |  |  |  |  |  |  |  |  |  |  |  |  |  |  |
| --- | --- | --- | --- | --- | --- | --- | --- | --- | --- | --- | --- | --- | --- | --- | --- | --- | --- | --- | --- | --- | --- | --- | --- | --- | --- | --- |
|  | CCP1<br>(43-104) |  |  |  |  |  |  |  |  |  |  |  |  | CCP2<br>(105-168) |  |  |  |  | CCP3<br>(169-234) |  |  |  | CCP4<br>(235-294) |  | CYT<br>(333-368) |  |
|  | 69 | 70 | 73 | 74 | 79 | 82 | 83 | 84 | 85 | 86 | 87 | 88 | 104 | 106 | 109 | 124 | 145 | 153 | 155 | 185 | 197 | 224 | 228 | 244 | 272 | 340 |
| NP_001229491.2 | V | Y | R | L | V | G | Q | V | L | A | L | V | K | R | T | S | N | S | T | N | T | E | Q | H | Y | R |
| XP_070624793.1 | . | . | . | . | I | . | . | . | . | . | . | I | . | . | . | . | . | . | . | . | A | . | . | . | . | . |
| MDBK | . | . | . | . | I | . | . | . | . | . | . | . | . | . | . | . | . | . | . | . | A | . | . | . | . | . |
| Zacarias | L | H | . | P | I | . | . | . | . | . | . | . | R | Q | N | N | . | . | . | S | . | . | K | Y | Q | G |
| Ginger | . | . | H | . | . | A | L | P | T | F | S | . | . | . | . | N | S | - | N | S | . | K | . | Y | Q | G |

To directly evaluate the functional capacity of the Zacarias *CD46* allele, we used a previously generated MDBK-CD46Δ cell line lacking endogenous *CD46* expression (8). Because MDBK cells have restricted lentiviral transduction efficiency, we further modified this *CD46*-null cell line by disrupting the bovine *TRIM5* retroviral restriction factor to enhance lentiviral delivery (**Fig. 4A**). Clones carrying edited *TRIM5* alleles were screened functionally using an eGFP-expressing lentivirus, and one clone showed a substantial increase in transduction efficiency compared with the parental population (approximately 34% versus 5%), consistent with reduced TRIM5-mediated restriction (**Fig. 4B**). Whole-genome sequence analyses showed that this clone was a compound heterozygote with an 8-nucleotide deletion on one allele and an in-frame 6-nucleotide deletion on the other (**Fig. 4C**). This MDBK-CD46Δ-TRIM5KD clone was subsequently used for lentiviral complementation studies.

**Figure 4.**
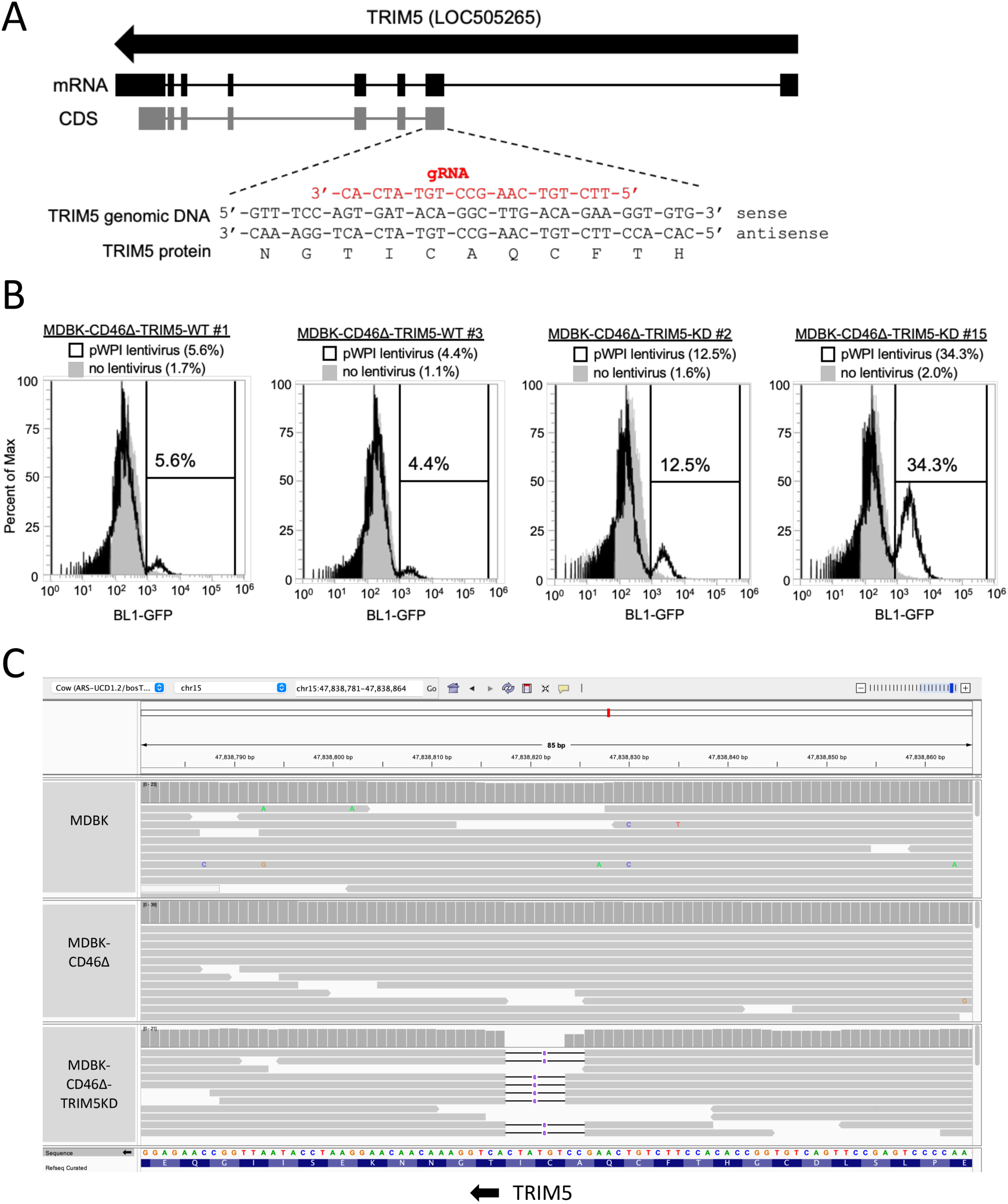
Generation of an MDBK-CD46Δ-TRIM5KD cell line. **A.** Map of bovine *TRIM5* gene with annotated guide RNA (gRNA). **B.** Flow cytometry analysis of eGFP expression to functionally screen individual MDBK-CD46Δ-TRIM5 clones for enhanced lentivirus transduction. Gated on percent cells expressing eGFP. **C.** Integrative Genomics Viewer (IGV) screenshot of the *TRIM5* exon 2 in MDBK-CD46Δ-TRIM5 clone 15, showing a compound heterozygous deletion.

Before generating the *CD46* complementation constructs, long-read RNA sequencing of naïve MDBK cells was used to identify the predominant endogenous *CD46* transcript isoform, which corresponded to the isoform represented by the bovine CD46 reference protein NP_001229491 but contained MDBK-specific sequence differences (**Table 2**). Native MDBK, wild-type Zacarias, and edited Ginger *CD46* constructs were generated using this same transcript isoform, allowing functional comparisons to focus on allele-specific sequence differences rather than differences in transcript isoform structure (**File S2**).

MDBK-CD46Δ-TRIM5KD cells were transduced with lentiviral vectors expressing each *CD46* construct under the constitutive EF1α promoter. Flow cytometric analysis showed that EF1α drove a highly homogeneous population with a narrow CD46 surface expression profile closely matching the distribution observed in parental MDBK cells (**Fig. 5A**). However, the overall surface expression was approximately two-fold greater in these transduced cells compared to endogenous levels observed in the parental MDBK cell line (**Fig. 5B**), with no significant difference in expression among the three transduced variants. Under these conditions, both the native MDBK and the wild-type Zacarias *CD46* alleles restored BVDV susceptibility in the *CD46* knockout cells. In contrast, expression of the Ginger allele failed to restore BVDV susceptibility, and these cells were not statistically different from the untransduced MDBK-CD46Δ-TRIM5KD control cells (**Fig. 5C**).

**Figure 5.**
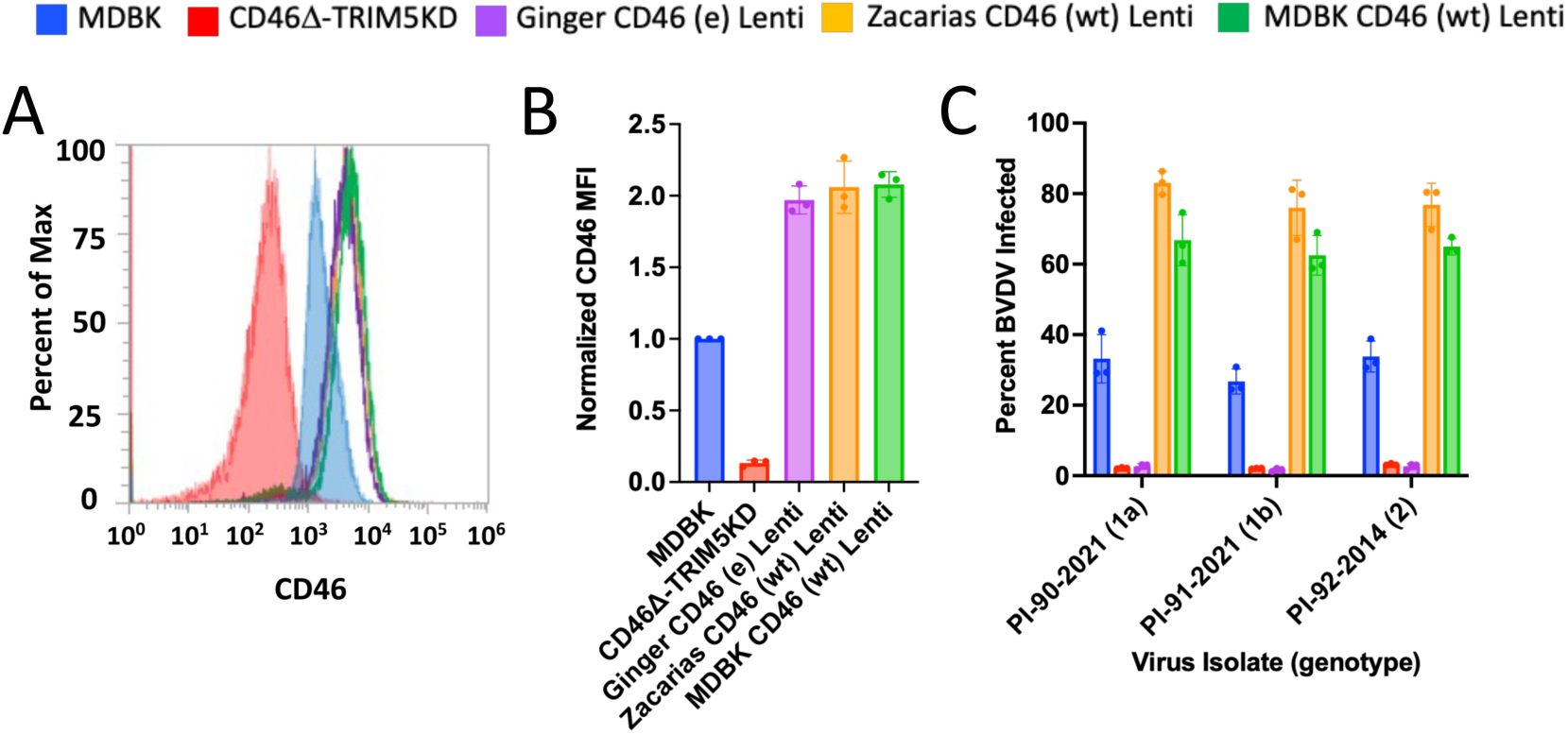
Functional analysis of EF1α-driven *CD46* variants in lentivirus complemented MDBK-CD46Δ-TRIM5KD cells. **(A)** Representative flow cytometric histogram overlay of EF1α-driven CD46 expression. The red filled histogram is MDBK-CD46Δ-TRIM5KD cells and the blue filled histogram is MDBK. The histograms for the lentivirus-complemented cells are unfilled. **(B)** Flow cytometric quantification of EF1α-driven CD46 surface expression. Surface expression levels (mean fluorescence intensity, MFI) are normalized to parental MDBK cells. Results are derived from three independent analysis dates using a single, stably transduced cell line per variant. **(C)** BVDV infection efficiency of EF1α-driven CD46 expression cell lines. Cells were infected with non-cytopathic (ncp) BVDV isolates belonging to genotypes 1a, 1b, or 2, and infection efficiency was determined at 20 hpi via flow cytometry using a monoclonal anti-BVDV E2 antibody. Data represent the mean ± standard deviation of three independent infection experiments using a single cell line per *CD46* variant. Abbreviations: e, edited *CD46* allele; wt, wild-type (unedited) *CD46* allele.

Given that the EF1α-driven expression exceeded endogenous CD46 protein levels, we next asked whether receptor overexpression might mask differences in receptor function or alter physiological BVDV entry. We therefore evaluated three alternative promoters, cytomegalovirus (CMV), spleen focus-forming virus (SFFV), and a chicken beta-actin (CAG), to identify one that more closely reproduced endogenous *CD46* expression levels. Among these, the SFFV promoter produced surface expression levels most similar to that of parental MDBK cells, averaging approximately 70% of endogenous expression (**Fig. 6B**). Unlike the EF1α promoter, however, SFFV generated broader distributions of CD46 surface expression (**Fig. 6A**). To account for this variability and minimize clonal bias, we established multiple independent stable cell lines for each *CD46* variant, including four expressing the native MDBK allele, three expressing the wild-type Zacarias allele, and two expressing the edited Ginger allele. Subsequent viral challenge of these SFFV-driven *CD46* variants confirmed that both the native MDBK and Zacarias alleles successfully restored BVDV permissiveness, even at lower surface densities (**Fig. 6C**). While minor variations in infection levels were observed between these two functional *CD46* alleles across different viral strains, both consistently supported BVDV infection. In contrast, expression of the Ginger allele did not restore susceptibility in the *CD46* knockout cells. These results demonstrate that the naturally occurring polymorphisms in the wild-type Zacarias *CD46* allele do not significantly impair receptor function, indicating that the reduced susceptibility of the heterozygous calf Giraldo is unlikely to result from impaired function of the paternal allele.

**Figure 6.**
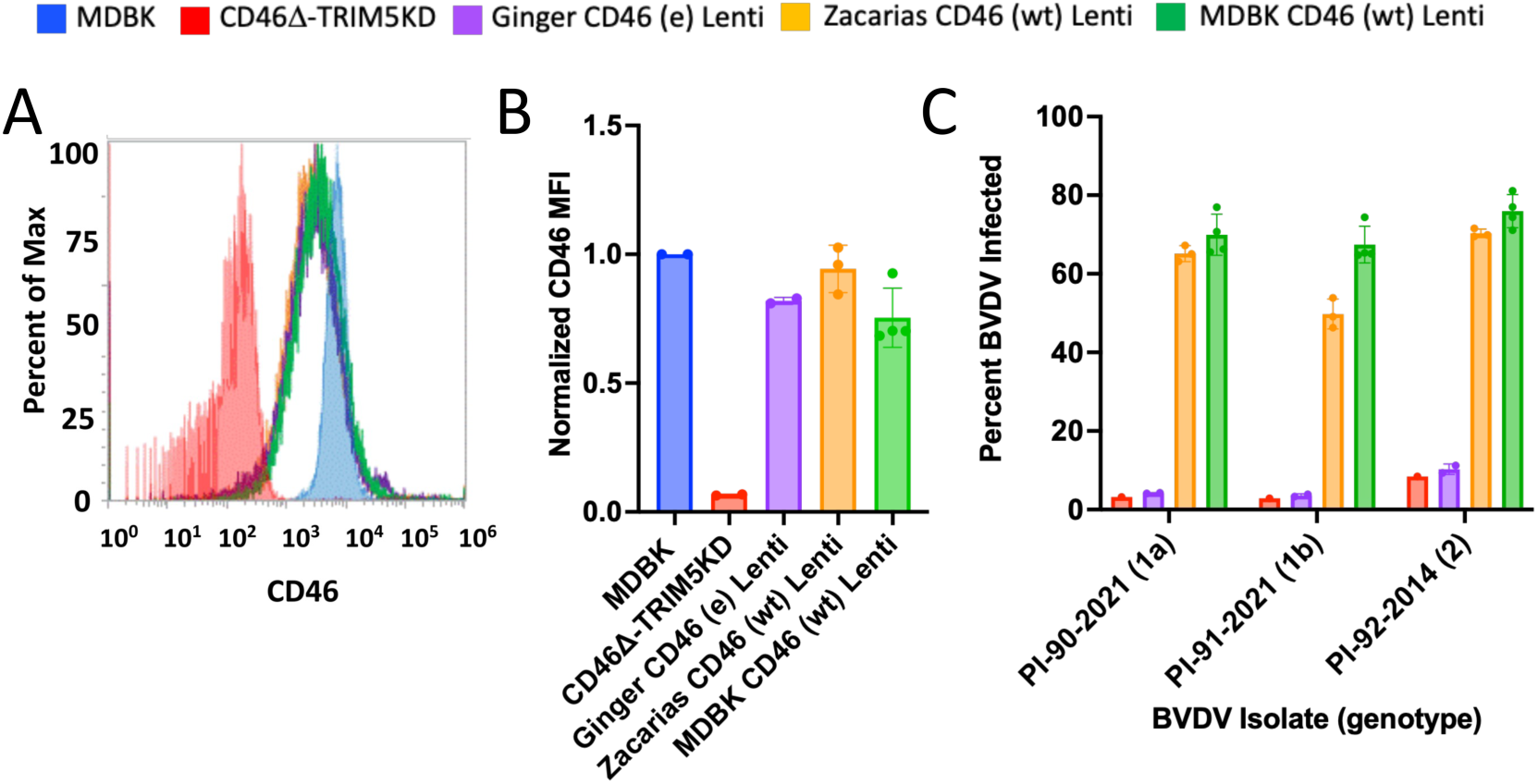
Functional analysis of SFFV-driven *CD46* variants in lentivirus complemented MDBK-CD46Δ-TRIM5KD cells. **(A)** Representative flow cytometric histogram overlay of MDBK cells compared to SFFV-driven CD46 expression. The red filled histogram is MDBK-CD46Δ-TRIM5KD cells and the blue filled histogram is MDBK. The histograms for the lentivirus-complemented cells are unfilled. **(B)** Flow cytometric quantification of SFFV-driven CD46 surface expression. CD46 protein surface expression levels are normalized to parental MDBK cells. Individual data points represent distinct, independently derived stable cell lines (native MDBK allele, n=4; Zacarias allele, n=3; Ginger allele, n=2). For each line, surface expression was measured in technical duplicate, averaged, and plotted as a single biological replicate to calculate the final group mean and standard deviation. **(C)** BVDV infection efficiency of SFFV-driven CD46 expression cell lines. Cells were challenged with ncp BVDV genotypes 1a, 1b, or 2, and infection rates were determined at 20 hpi. Individual data points represent the identical set of independent cell lines described in panel B. BVDV infections were executed in technical duplicate, averaged, and plotted as a single biological replicate to determine the group mean and standard deviation across cell lines to account for expression heterogeneity and reduce clonal bias. Abbreviations: e, edited *CD46* allele; wt, wild-type (unedited) *CD46* allele.

## Discussion

This study provides evidence that inheritance of a single *CD46* A_82_LPTFS allele is associated with reduced BVDV susceptibility in a heterozygous animal in *ex vivo* assays. Previous work established reduced BVDV susceptibility in a homozygous edited heifer (8); however, whether a single edited *CD46* allele could produce a comparable phenotype remained unknown. Here, the Gir bull calf, Giraldo, inherited a single edited *CD46* allele and exhibited reduced *ex vivo* BVDV susceptibility across multiple primary cell types at levels comparable to those observed in his homozygous edited dam, Ginger. This phenotype was observed against representative isolates from two BVDV genotypes and biotypes, including non-cell culture-adapted isolates obtained from persistently infected cattle. Although these findings from a single heterozygous animal are consistent with a dominant effect of the edited allele, additional heterozygous animals will be required to determine whether inheritance of a single edited allele consistently results in reduced BVDV susceptibility. Likewise, confirmation through future *in vivo* BVDV challenge studies will be necessary to determine whether *ex vivo* reductions in susceptibility translate to protection at the animal level. If confirmed, this inheritance pattern could enable more rapid dissemination of reduced BVDV susceptibility through conventional cattle breeding strategies.

To date, most livestock disease-resistance edits that have been examined in heterozygous animals have shown that a single modified allele is insufficient to confer resistance, with protection generally requiring biallelic modification. For example, pigs carrying a homozygous deletion of the SRCR5 domain of *CD163* are resistant to porcine reproductive and respiratory syndrome virus (PRRSV), whereas heterozygous animals remain susceptible (17). Similarly, homozygous editing of the avian leukosis virus subgroup J (ALV-J) receptor *chNHE1* confers resistance in chickens, while heterozygous birds remain susceptible (18). Other host-directed strategies, including modification of *DNAJC14* to limit pestivirus replication in pigs (19), modification of *ANP32A* to reduce avian influenza virus (AIV) polymerase activity in chickens (20), and *ANPEP* disruption to reduce transmissible gastroenteritis virus (TGEV) susceptibility in pigs (21), have primarily been evaluated in homozygous edited animals, with comparatively little characterization of the effects of heterozygosity. Similar patterns are observed among naturally occurring host variants that influence viral susceptibility. For example, resistance-associated variants in the human *CCR5* and *FUT2* genes confer resistance to HIV-1 and several norovirus strains, respectively, primarily in homozygous individuals, whereas heterozygous individuals generally remain susceptible to infection (22–24). Although some heterozygous variants can influence disease progression, substantial reductions in viral susceptibility mediated by a single altered host allele appear uncommon.

Unlike receptor knockout strategies, the *CD46* A_82_LPTFS edit modifies the viral interaction surface without eliminating receptor expression. The observation that a heterozygous animal nevertheless displays reduced *ex vivo* BVDV susceptibility despite continued expression of a wild-type *CD46* allele raises important questions regarding the molecular basis of this phenotype. One possibility is that the edited CD46 protein interferes with wild-type receptor function in a dominant-negative-like manner, whereas an alternative explanation is a threshold effect resulting from reduced functional receptor abundance. The dominant-negative model aligns with the entry mechanisms of other CD46-utilizing viruses, such as measles virus, which rely on multivalent interactions and receptor clustering to initiate endocytosis (25). The edited CD46 protein could participate in non-productive associations with wild-type CD46 protein, reducing the pool of receptors available for productive clustering. However, key mechanistic details remain unresolved. It is not yet known whether BVDV binding induces spatial reorganization of CD46, and the native oligomerization state of bovine CD46 remains undefined (26). In the absence of these data, a simpler threshold-based model cannot be excluded, in which the reduced abundance of functional wild-type receptors in heterozygotes falls below a critical density required for infection. Distinguishing between these models will require direct assessment of CD46 organization and receptor function during BVDV attachment and entry.

Additionally, future allele-specific disruption of the edited *CD46* allele in Giraldo’s heterozygous cells will help determine whether reduced BVDV susceptibility requires the presence of the edited protein or is primarily attributable to reduced functional CD46 receptor abundance. Although reduced BVDV susceptibility was observed across all isolates tested, the magnitude of protection varied among the virus isolates. These isolate-dependent differences in residual infectivity may reflect variation in CD46 dependence or the ability of individual viruses to use alternative entry mechanisms. It is well-documented that BVDV can adapt during cell culture propagation, leading to altered receptor usage and viral entry properties (27). Here, both BVDV-2 isolates exhibited relatively greater infectivity in edited lymphocytes than the nine BVDV-1 isolates examined. This apparent difference was less pronounced in our previous study using an earlier passage of the same BVDV-2 isolates, suggesting that *in vitro* adaptation may have contributed to altered receptor usage or entry properties (8, 27). Because primary bovine cells completely lacking CD46 were not available, we could not determine whether residual infection of lymphocytes reflected CD46-independent entry or isolate-specific differences in the ability to engage the edited CD46 receptor. Thus, the biological and *in vivo* significance of BVDV-2 infectivity in *CD46*-edited cells remains to be determined during natural infection.

The translational success of gene-editing technology also depends on animal fitness. CD46 has been implicated in mammalian fertilization (14). However, Ginger exhibited normal oocyte fertilization during *in vitro* fertilization (15), conceived following artificial insemination, carried a pregnancy to term, and produced a healthy offspring. These findings suggest that the *CD46* edit does not impair female reproductive function under the conditions evaluated. While Giraldo appears healthy and developmentally normal, he has not yet reached sexual maturity, precluding evaluation of male reproductive function. Thus, the fertility of *CD46*-edited males and the long-term fitness of edited animals across multiple generations remain important outstanding questions.

In summary, the finding that a single edited *CD46* allele is associated with significantly reduced BVDV susceptibility expands the practical potential of this approach. If this phenotype is confirmed in additional heterozygous animals through *in vivo* challenge studies, this trait could be incorporated into existing artificial insemination and conventional breeding programs using homozygous edited sires. These findings support the broader application of precise genome editing to modify host-pathogen interactions and improve disease resilience in livestock.

## Materials and Methods

### Animal Ethics Statement

All animal procedures were approved by the University of Nebraska–Lincoln Institutional Animal Care and Use Committee (IACUC; Projects 2711 and 2466).

### Reproductive Procedures

Ginger was synchronized for estrus using a 7-day Controlled Internal Drug Release (CIDR)-based Controlled Ovulation Synchronization (CO-Synch) protocol (28) and artificially inseminated with semen from two unedited Gir sires. Pregnancy was confirmed by ultrasound as previously described (15).

### Whole Genome Sequence (WGS) Analysis for Paternity and Editing

WGS was performed with whole blood from Giraldo and semen from the two potential sires: Zacarias De Brasuca FIV, Brazil 674021, 12/11/2011, American Brahman Breeders Association (ABBA) registration number 955764 (STgenetics, Navasota, TX); and Zamuel De Brasuca FIV, Brazil 699544, 5/7/2012, ABBA 955768 (STgenetics). DNA was extracted using standard procedures (29) and 500 bp paired-end libraries were prepared as previously described (8). Reads were mapped to ARS-UCD1.2 and aligned genomes were viewed with the Integrative Genomics Viewer (IGV) version 2.12.2. Paternity was assigned by allelic exclusion using 115 previously validated parentage SNPs (**Table S1**), which distinguished Zacarias from the alternative potential sire. *CD46* allele inheritance was confirmed by inspection of the relevant genomic region. Detailed sequencing and parentage-analysis procedures are provided in Supplemental Methods.

MDBK-CD46Δ-TRIM5KD cells used in lentivirus transduction studies were also sequenced (see “TRIM5 Knockdown in MDBK-CD46Δ cells” section for details). The WGS for the parental MDBK cell line and the unedited TO470 Gir cell line were previously published (MDBK: (30); TO470: (8)). The TO470 cell line is an unedited Gir breed female fibroblast line that served as the source for the gene-editing and subsequent somatic cell nuclear transfer (SCNT) to produce Ginger (8).

### Cell Lines and Culture Conditions

BVDV-free Madin-Darby bovine kidney (MDBK) cells (ATCC CCL-22, Lot no. 375272) were obtained from the American Type Culture Collection (ATCC; Rockville, MD, USA). A *CD46* gene-deleted derivative of this line, designated as MDBK-CD46Δ, was generated previously (8). Both cell lines were cultured in minimum essential medium (MEM) (Gibco, Grand Island, NY, USA) supplemented with 10% gamma-irradiated fetal bovine serum (FBS) (Atlas Biologicals, Fort Collins, CO, USA), 1× antibiotic–antimycotic (Gibco, Grand Island, NY, USA), and 2 mM L-Glutamine (Gibco). 293T cells (ATCC CRL-3216) were propagated in DMEM (Corning, Tewksbury, MA, USA), supplemented with 10% bovine serum (Peak Serum, Bradenton, FL, USA) and 1× antibiotic–antimycotic (Gibco). Primary skin fibroblasts from the *CD46* heterozygous calf (Giraldo) were isolated from an ear punch tissue biopsy (Allflex tissue sampling unit) at Trans Ova Genetics (Sioux Center, IA, USA). Primary fibroblasts for the unedited Gir line (TO470), the homozygous *CD46*-edited Gir dam (Ginger), and a wild-type unedited Holstein control (Mary Ann) were previously established (8). The primary fibroblasts were cultured in DMEM (Corning) supplemented with 15% irradiated FBS (Atlas Biologicals) and 1× antibiotic–antimycotic (Gibco).

### Virus Isolates and Serum from Calves Persistently Infected with BVDV

BVDV-1 and BVDV-2 isolates were obtained from serum collected from persistently infected cattle between 2014 and 2023. Virus was isolated and propagated in MDBK cells, and titers of noncytopathic isolates were determined by flow cytometry-based endpoint dilution using an anti-BVDV E2 antibody (VMRD, Pullman, WA, USA) as previously described (8, 27).

### Flow Cytometric Quantification of CD46 Surface Expression and BVDV Infection

CD46 surface expression and BVDV infection were quantified by flow cytometry as previously described (8, 27). Briefly, fixed cells were stained with a custom anti-bovine CD46 antibody or anti-BVDV E2 antibody (VMRD, Catalog no. 348 for MDBK and primary fibroblasts and Creative Diagnostics, Shirley, NY, USA, Catalog no. DMAB28412 for lymphocytes) followed by fluorescent secondary antibodies and analyzed using an Attune NxT flow cytometer (Thermo Fisher Scientific, Waltham, MA, USA). Detailed procedures are provided in Supplemental Methods.

### Immunofluorescence Microscopy

CD46 protein localization was assessed by immunofluorescence as previously described (8), using a custom anti-bovine CD46 antibody, a FITC-conjugated secondary antibody, and DAPI nuclear counterstaining (8).

### BVDV Susceptibility Assays in Primary and Immortalized Lentivirus Transduced Cell Lines

Susceptibility to BVDV was evaluated in both primary skin fibroblasts and MDBK-CD46Δ-TRIM5KD cells expressing various *CD46* variants (see the “TRIM5 Knockdown in MDBK-CD46Δ cells” section for details on cell line creation). Cells were seeded in 24-well plates at a density calculated to achieve approximately 80% confluency the following day. To measure CD46-dependent virus entry, BVDV isolates were pre-incubated with 200 µg/mL heparin (H3149-10KU, Sigma-Aldrich, St. Louis, MO, USA) in serum-free medium for 30 min at 37°C. Heparin blocks the heparan sulfate binding sites on BVDV, thereby blocking a common alternative entry pathway for some BVDV strains (27). Cells were inoculated with noncytopathic BVDV isolates at a multiplicity of infection (MOI) empirically determined to achieve a consistent infection rate in susceptible control cells (approximately 80%). Following a 2-hour incubation at 37°C for virus adsorption and entry, unbound virus was removed by washing. Cells were then incubated for 20 hours and BVDV infection was quantified by flow cytometric detection of the viral E2 glycoprotein, as described above. To test non-cell culture-adapted BVDV, serum collected from cattle persistently infected with BVDV was directly inoculated onto cells. Following a 2-hour incubation, cells were washed and incubated for 72 hours.

### Monocyte and Lymphocyte Isolation and *Ex Vivo* BVDV Challenge

Peripheral blood mononuclear cells (PBMCs) were isolated from Ginger, Giraldo, and the age-and sex-matched control calf Bueno by density-gradient centrifugation using SepMate tubes. Monocytes and lymphocytes were isolated as previously described (8). For lymphocyte susceptibility assays, 2.5×10^5^ cells were inoculated with 80 µL of BVDV inoculum per well in 96-well round-bottom plates, corresponding to an average MOI of approximately 5 across isolates. Infection was quantified at 32 hpi by flow cytometric detection of intracellular BVDV E2 antigen (8). For monocyte susceptibility assays, 4×10^5^ cells were challenged with BVDV isolates at an MOI of 0.01 or with a 1:10 dilution of serum collected from persistently infected cattle. Viral replication was quantified at 48 hpi for serum challenges and 72 hpi for isolate challenges by BVDV-specific RT-qPCR (31), and viral RNA was expressed as fold change relative to the input control. Detailed cell isolation, inoculation, incubation, and RT-qPCR conditions are provided in Supplemental Methods.

### Long-Read Transcript Analysis of *CD46* Allele Expression and Isoform Selection

Long-read transcript sequencing was used to assess *CD46* allele expression in Giraldo-derived cells and identify the predominant *CD46* transcript isoform for the lentiviral complementation model. Full-length mRNA libraries from whole blood and primary skin fibroblasts were prepared using the PacBio Iso-Seq/Kinnex workflow and sequenced on the Revio platform according to the manufacturer’s protocol (Pacific Biosciences, Menlo Park, CA, USA).

Full-length non-concatemer (FLNC) reads were screened for allele-specific sequence variation corresponding to the wild-type G_82_QVLAL or edited A_82_LPTFS *CD46* alleles. Custom scripts were used to count transcripts containing each allele-specific sequence and estimate relative allele representation (Supplemental Methods).

A separate analysis was performed to identify the predominant *CD46* transcript isoform expressed in MDBK cells for use in *CD46* complementation constructs. Previously generated PacBio Iso-Seq data from MDBK cells were reanalyzed for this purpose. Total RNA from MDBK cells was sequenced using the PacBio Iso-Seq platform in April 2021, and raw sequencing data were processed by the USMARC Core Lab using the PacBio Iso-Seq analysis pipeline to generate collapsed consensus transcript isoforms and associated full-length read counts. Consensus transcript sequences were imported into Geneious Prime and aligned to RefSeq accession no. NM_001242561 to identify the corresponding *CD46* transcript cluster. Isoform abundance was then determined based on the number of supporting full-length reads (count_fl) reported by the Iso-Seq pipeline.

### Short-Read mRNA Expression Analysis of *CD46*

Short-read stranded mRNA libraries (TruSeq, Illumina, San Diego, CA, USA) were prepared from the same RNA samples used for long-read mRNA sequencing to provide an independent measure of relative allele expression. Paired-end short reads (2×150 bp) were sequenced with a high-throughput benchtop sequencer and sequenced by synthesis chemistry (NextSeq2000, Illumina). Raw sequence reads were trimmed using the BBDuk plugin (32) within Geneious Prime (version 2026.1.1) to remove sequence adapters and low-quality bases. Trimmed and filtered reads were then aligned to exons 1-6 of the two *CD46* alleles sequenced in the *CD46* heterozygous calf (Giraldo) with the native Geneious alignment algorithm. Mapped reads encoding the wild-type G_82_QVLAL or the edited A_82_LPTFS sequence were counted to provide an independent measure of relative allele expression.

### *TRIM5* Knockdown in MDBK-CD46Δ Cells

MDBK-CD46Δ cells were modified by CRISPR/Cas9-mediated non-homologous end joining (NHEJ) targeting of bovine TRIM5-3 to enhance lentiviral transduction (33, 34). Edited clones were isolated by single-cell sorting and screened by amplicon sequencing. A clone exhibiting increased lentiviral transduction was characterized by whole genome sequence and selected for subsequent complementation studies. Detailed editing and genotyping procedures are provided in Supplemental Methods.

### Lentivirus Production

Lentiviral vectors expressing the indicated *CD46* variants under constitutive EF1α (pTwist Lenti EF1α EGFP Puro) or SFFV (pTwist Lenti SFFV Puro WPRE) promoters were generated in 293T cells using a third-generation packaging system (pMD2.G, pRSV-REV, and pMDLg/pRRE). CD46 constructs used the same predominant bovine *CD46* transcript isoform and were sequence verified. Lentiviral supernatants were concentrated by centrifugal filtration and used to transduce MDBK-CD46Δ-TRIM5KD cells. Transduced cells were selected with puromycin and subsequently used for CD46 expression and BVDV susceptibility assays. Detailed vector, lentivirus production, transduction, and selection procedures are provided in Supplemental Methods.

## Statistical Analyses

Statistical analyses were performed using GraphPad Prism (version 10.2.3) (GraphPad Software, LLC, Boston, MA, USA). Statistical significance was defined as *p* < 0.05. For CD46 protein expression, a one-way analysis of variance (ANOVA) was used to compare CD46 expression among cell lines, followed by Tukey’s multiple-comparisons test when the overall ANOVA was significant. For BVDV susceptibility assays with complete data across experimental dates, two-way ANOVA was used. For assays containing missing values due to differences in cell availability among experimental dates, two-way mixed-effects models were used. In both analyses, BVDV isolate and animal/cell source were included as factors, and the full model was fitted. Šídák’s multiple-comparisons test was used for pairwise comparisons among animal/cell sources within individual BVDV isolates. For allele-specific CD46 expression analyses, deviations from the expected 1:1 allelic ratio were assessed separately for each sample and sequencing approach using Pearson’s chi-square goodness-of-fit test.

## Data Availability

The genomic sequence data have been deposited with links to BioProject accession number PRJNA887820 in the NCBI BioProject database with the corresponding sequencing datasets available through the associated NCBI Sequence Read Archive (SRA) records. Other data generated during this study are available from the corresponding author upon reasonable request.

## Acknowledgements

We thank Susan Hauver, Nathan Allison, and the USMARC Core Facility for technical support, the University of Nebraska animal care staff, and Donna Griess for secretarial and administrative support. Funding for this research was provided by the USDA, ARS appropriated project 3040-32000-036-00D (A.M.W. and M.P.H.). The authors declare that all scientific content, analyses, interpretations, and conclusions were developed and verified by the authors, who take full responsibility for the integrity, accuracy, and originality of the work. Artificial intelligence tools (Gemini, ChatGPT, and Claude) were used during manuscript preparation for language editing, including improvement of grammar, clarity, and conciseness. AI tools were not used to generate scientific content, data interpretations, figures, tables, code, or conclusions. Mention of trade names or commercial products in this publication is solely for the purpose of providing specific information and does not imply recommendation or endorsement by the U.S. Department of Agriculture. The USDA is an equal opportunity provider and employer.

## Author Contributions

A.M.W., A.C.K., M.P.H., A.P.S., T.S., and B.L.V.L. conceived and designed the study. A.M.W., A.C.K., M.P.H., A.P.S., K.K., and B.L.V.L. performed the experiments. A.M.W., A.C.K., and M.P.H. analyzed and interpreted the data. T.S. provided animals, and K.K. provided US MARC core laboratory expertise and reagents. A.M.W., M.P.H., T.S., and B.L.V.L. supervised the study. A.M.W. wrote the initial manuscript draft, and all authors reviewed and edited the manuscript. All authors approved the final manuscript.

## Supplementary Materials

Supplemental Methods

Table S1. Bovine SNP list for 115 highly informative Parentage SNPs

File S2. *CD46* open reading frame sequences used for lentivirus complementation studies

## References

1. Walz PH, Chamorro MF, Falkenberg SM, Passler T, van der Meer F, Woolums AR. 2020. Bovine viral diarrhea virus: An updated American College of Veterinary Internal Medicine consensus statement with focus on virus biology, hosts, immunosuppression, and vaccination. J Vet Intern Med 34:1690–1706.

2. Richter V, Lebl K, Baumgartner W, Obritzhauser W, Kasbohrer A, Pinior B. 2017. A systematic worldwide review of the direct monetary losses in cattle due to bovine viral diarrhoea virus infection. Vet J 220:80–87.

3. Houe H. 2003. Economic impact of BVDV infection in dairies. Biologicals 31:137–43.

4. McClurkin AW, Littledike ET, Cutlip RC, Frank GH, Coria MF, Bolin SR. 1984. Production of cattle immunotolerant to bovine viral diarrhea virus. Can J Comp Med 48:156–61.

5. Coggins L, Gillespie JH, Robson DS, Thompson JD, Phillips WV, Wagner WC, Baker JA. 1961. Attenuation of virus diarrhea virus (Strain Oregon C24V) for vaccine purposes. Cornell Vet 51:539–45.

6. van Oirschot J, Bruschke CJM, van Rijn PA. 1999. Vaccination of cattle against bovine viral diarrhoea. Vet Microbiol 64:169–183.

7. Maurer K, Krey T, Moennig V, Thiel HJ, Rumenapf T. 2004. CD46 is a cellular receptor for bovine viral diarrhea virus. J Virol 78:1792–9.

8. Workman AM, Heaton MP, Vander Ley BL, Webster DA, Sherry L, Bostrom JR, Larson S, Kalbfleisch TS, Harhay GP, Jobman EE, Carlson DF, Sonstegard TS. 2023. First gene-edited calf with reduced susceptibility to a major viral pathogen. PNAS Nexus 2:pgad125.

9. Krey T, Himmelreich A, Heimann M, Menge C, Thiel HJ, Maurer K, Rumenapf T. 2006. Function of bovine CD46 as a cellular receptor for bovine viral diarrhea virus is determined by complement control protein 1. J Virol 80:3912–22.

10. Workman AM, Heaton MP, Vander Ley BL. 2025. CD46 gene editing confers ex vivo BVDV resistance in fibroblasts from cloned Angus calves. Viruses 17:775.

11. Liszewski MK, Kemper C. 2019. Complement in motion: The evolution of CD46 from a complement regulator to an orchestrator of normal cell physiology. J Immunol 203:3–5.

12. Yamamoto H, Fara AF, Dasgupta P, Kemper C. 2013. CD46: the ’multitasker’ of complement proteins. Int J Biochem Cell Biol 45:2808–20.

13. Jankovicova J, Antalikova J, Simon M, Michalkova K, Horovska L. 2011. Comparative fluorescence analysis of the bovine sperm using IVA-520 (anti-CD46 antibody) and lectins: probable localisation of CD46 on bovine sperm membrane. Gen Physiol Biophys 30 Spec No:S70-6.

14. Liszewski MK, Atkinson JP. 2021. Membrane cofactor protein (MCP; CD46): deficiency states and pathogen connections. Curr Opin Immunol 72:126–134.

15. Snider AP, Workman AM, Heaton MP, Vander Ley BL, Krueger AC, Sonstegard TS. 2025. Fertility and early embryonic development in a *CD46*-edited Gir heifer with reduced susceptibility to BVDV. Biol Reprod 112:245–252.

16. Zezafoun H, Decreux A, Desmecht D. 2011. Genetic and splice variations of Bos taurus CD46 shift cell permissivity to BVDV, the bovine pestivirus. Vet Microbiol 152:315–27.

17. Burkard C, Lillico SG, Reid E, Jackson B, Mileham AJ, Ait-Ali T, Whitelaw CBA, Archibald AL. 2017. Precision engineering for PRRSV resistance in pigs: Macrophages from genome edited pigs lacking CD163 SRCR5 domain are fully resistant to both PRRSV genotypes while maintaining biological function. PLoS Pathog 13:e1006206.

18. Koslová A, Trefil P, Mucksová J, Reinisová M, Plachy J, Kalina J, Kucerová D, Geryk J, Krchlíková V, Lejcková B, Hejnar J. 2020. Precise CRISPR/Cas9 editing of the NHE1 gene renders chickens resistant to the J subgroup of avian leukosis virus. Proc Natl Acad Sci U S A 117:2108–2112.

19. Crooke HR, Schwindt S, Fletcher SL, Isken O, Harding S, Berkley N, Tait-Burkard C, Warren C, Whitelaw CBA, Tautz N, Lillico SG. 2026. DNAJC14 gene-edited pigs are resistant to classical pestiviruses. Trends Biotechnol 44:570–586.

20. Idoko-Akoh A, Goldhill DH, Sheppard CM, Bialy D, Quantrill JL, Sukhova K, Brown JC, Richardson S, Campbell C, Taylor L, Sherman A, Nazki S, Long JS, Skinner MA, Shelton H, Sang HM, Barclay WS, McGrew MJ. 2023. Creating resistance to avian influenza infection through genome editing of the ANP32 gene family. Nat Commun 14:6136.

21. Whitworth KM, Rowland RRR, Petrovan V, Sheahan M, Cino-Ozuna AG, Fang Y, Hesse R, Mileham A, Samuel MS, Wells KD, Prather RS. 2019. Resistance to coronavirus infection in amino peptidase N-deficient pigs. Transgenic Res 28:21–32.

22. Dean M, Carrington M, Winkler C, Huttley GA, Smith MW, Allikmets R, Goedert JJ, Buchbinder SP, Vittinghoff E, Gomperts E, Donfield S, Vlahov D, Kaslow R, Saah A, Rinaldo C, Detels R, O’Brien SJ. 1996. Genetic restriction of HIV-1 infection and progression to AIDS by a deletion allele of the CKR5 structural gene. Science 273:1856–1862.

23. Lindesmith L, Moe C, Marionneau S, Ruvoen N, Jiang X, Lindblad L, Stewart P, LePendu J, Baric R. 2003. Human susceptibility and resistance to Norwalk virus infection. Nat Med 9:548–53.

24. Thorven M, Grahn A, Hedlund KO, Johansson H, Wahlfrid C, Larson G, Svensson L. 2005. A homozygous nonsense mutation (428G→A) in the human secretor (FUT2) gene provides resistance to symptomatic norovirus (GGII) infections. J Virol 79:15351–15355.

25. Santiago C, Celma ML, Stehle T, Casasnovas JM. 2010. Structure of the measles virus hemagglutinin bound to the CD46 receptor. Nat Struct Mol Biol 17:124–9.

26. Aitkenhead H, Stuart DI, El Omari K. 2023. Structure of bovine CD46 ectodomain. Viruses 15:1424.

27. Krueger AC, Vander Ley BL, Heaton MP, Sonstegard TS, Workman AM. 2025. Primary cells from a CD46-edited bovine heifer have reduced BVDV susceptibility despite viral adaptation to heparan sulfate. Viruses 17:634.

28. Hill SL, Perry GA, Mercadante VRG, Lamb GC, Jaeger JR, Olson KC, Stevenson JS. 2014. Altered progesterone concentrations by hormonal manipulations before a fixed-time artificial insemination CO-Synch plus CIDR program in suckled beef cows. Theriogenology 82:104–113.

29. Heaton MP, Grosse WM, Kappes SM, Keele JW, Chitko-McKown CG, Cundiff LV, Braun A, Little DP, Laegreid WW. 2001. Estimation of DNA sequence diversity in bovine cytokine genes. Mamm Genome 12:32–7.

30. Workman AM, Heaton MP, Webster DA, Harhay GP, Kalbfleisch TS, Smith TPL, Falkenberg SM, Carlson DF, Sonstegard TS. 2021. Evaluating large spontaneous deletions in a bovine cell line selected for bovine viral diarrhea virus resistance. Viruses 13:2147.

31. Mahlum CE, Haugerud S, Shivers JL, Rossow KD, Goyal SM, Collins JE, Faaberg KS. 2002. Detection of bovine viral diarrhea virus by TaqMan reverse transcription polymerase chain reaction. J Vet Diagn Invest 14:120–5.

32. Bushnell B, Rood J, Singer E. 2017. BBMerge -Accurate paired shotgun read merging via overlap. PLoS One 12:e0185056.

33. Si Z, Vandegraaff N, O’Huigin C, Song B, Yuan W, Xu C, Perron M, Li X, Marasco WA, Engelman A, Dean M, Sodroski J. 2006. Evolution of a cytoplasmic tripartite motif (TRIM) protein in cows that restricts retroviral infection. Proc Natl Acad Sci U S A 103:7454–9.

34. Sawyer SL, Emerman M, Malik HS. 2007. Discordant evolution of the adjacent antiretroviral genes TRIM22 and TRIM5 in mammals. PLoS Pathog 3:e197.

