## Supplemental Files for "Inheritance of a Single Edited *CD46* Allele Is Associated with Reduced *Ex Vivo* Susceptibility to Bovine Viral Diarrhea Virus"

1. Supplemental Methods

2. Table S1. Bovine SNP List for 115 Highly Informative Parentage SNPs

3. File S2. *CD46* Open Reading Frame Sequences used for Lentivirus Complementation Studies

### Supplemental Methods

#### Whole Genome Sequence (WGS) Analysis for Paternity and Editing

To confirm the paternity and genotype conformation of Giraldo, WGS was performed with whole blood from Giraldo and semen from the two potential sires: Zacarias De Brasuca FIV, Brazil 674021, 12/11/2011, American Brahman Breeders Association (ABBA) registration number 955764 (STgenetics, Navasota, TX, USA); and Zamuel De Brasuca FIV, Brazil 699544, 5/7/2012, ABBA 955768 (STgenetics). Unless otherwise indicated, all reagents were molecular-biology grade. The DNA from semen, blood, or cell lines was extracted using standard procedures that included RNase and protease digestion, phenol/chloroform extraction, and ethanol precipitation (1). Purified DNA was dissolved in a solution of 10 mM TrisCl, 1 mM EDTA (TE, pH 8.0) and stored at 4 °C.

For WGS, 1 µg of genomic DNA was fragmented and used to create indexed, 500 bp, paired-end libraries as previously described (2). Briefly, dynamic pools of indexed libraries were sequenced on massively parallel sequencing machines (NextSeq2000, Illumina Inc., San Diego, CA, USA) with appropriate manufacturer's kits to produce 2 × 150 bp paired-end reads. Pooled samples were sequenced until a minimum of 40-45 Gb of data was collected with a quality score of greater than Q20 for each animal, thereby producing at least 13- to 15-fold mapped read coverage for each animal (reference genome ARS-UCD1.2). This level of coverage provides scoring rates and SNP genotype accuracies that exceed 99% (3). The BAM files produced at each step were indexed with SAMtools and made available via object storage through a web service interface (Amazon Web Services, Inc. Seattle, WA). Aligned genomes were viewed with the Integrative Genomics Viewer (IGV) version 2.12.2 by selecting the desired reference genome and loading a session file URL. Inheritance of *CD46* alleles was scored by viewing the relevant regions in IGV.

Giraldo's paternity was established by allelic exclusion. Parentage exclusion is based on the principle that a parent and offspring must share an allele at every locus (4). Success of this approach requires high genotyping accuracy (≥99%), high minor allele frequency (MAF, ≥0.30), and a 10- to 30-Mb spacing across the 2.7 Gb genome (90 to 270 SNPs). Here, we used a set of 115 reference "parentage" SNPs (**Table S1**) that are also present on the Bovine HD 770 BeadArray (Illumina, San Diego CA). The IGV was used to visually score genotypes at these 115 sites for allele exclusion. The eight relevant sites from **Table S1** that yielded sire allelic exclusions were: chr1:29963400, chr1:149646917, chr5:27670995, chr10:97152695, chr21:26120612, chr22:55883157, chr27:37824983, chr28:5874287. These sites, together with the rest of the genome sites, can be viewed by loading this public session file URL directly in to IGV: https://s3.us-west-2.amazonaws.com/usmarc.heaton.public/WGS/CellLines/ARS1.2/sessions/ARS1.2_CD46_GingerGerryZacZam4Tracks.xml

In addition to sequencing the above samples, MDBK-CD46Δ-TRIM5KD cells used in lentivirus transduction studies were also sequenced (see "TRIM5 Knockdown in MDBK-CD46Δ cells" section for details). The WGS for the parental MDBK cell line and the unedited TO470 Gir cell line were previously published (MDBK: (5); TO470: (2)). The TO470 cell line is an unedited Gir breed female fibroblast line that served as the source for the gene-editing and subsequent somatic cell nuclear transfer (SCNT) to produce Ginger (2).

**Flow Cytometric Quantification of CD46 Surface Expression and BVDV Infection**

CD46 protein expression and BVDV infection were quantified using flow cytometry. Cells were harvested from culture plates by enzymatic dissociation with a proprietary trypsin-based reagent (TrypLE Express, Gibco, Grand Island, NY, USA) and collected by centrifugation. The cell pellet was washed three times with phosphate-buffered saline (PBS) and then fixed with 4% paraformaldehyde for 12 min at room temperature. After fixation, the cells were washed again with PBS and prepared for staining.

Cells were stained for CD46 by first blocking with 2% bovine serum albumin (BSA) in phosphate-buffered saline (PBS) for 30 min at room temperature. Cells were then incubated with a custom polyclonal rabbit anti-bovine CD46 antibody at a concentration of 1 µg/mL for 30 min (2). This was followed by incubation for 30 min in the dark with either a goat anti-rabbit IgG FITC-conjugated secondary antibody (Abcam, Cambridge, MA, USA, Cat. no. ab6717) or, for EF1α-driven CD46-expressing cells, a mouse anti-rabbit IgG PE-Cy7-conjugated secondary antibody (Santa Cruz Biotechnology, Dallas, TX, USA, Cat. no. 516721). Cells were washed three times with PBS and resuspended in 1 mL of PBS before flow cytometric analysis (Attune

NxT, Thermo Fisher Scientific, Waltham, MA, USA). Specificity controls, including cells lacking CD46, previously demonstrated the specificity of this custom CD46 antibody (2).

BVDV infection was detected in cells by first washing and blocking with a blocking and permeabilization buffer (PBS containing 2% BSA and 0.1% w/v saponin) for 30 min. The cells were then resuspended in 100 µL of an antibody dilution buffer (PBS containing 1% BSA and 0.1% w/v saponin) containing a 1:200 dilution of an anti-BVDV E2 monoclonal antibody (VMRD, Pullman, WA, USA, Catalog no. 348 for MDBK and primary fibroblasts and Creative Diagnostics, Shirley, NY, USA, Catalog no. DMAB28412 for lymphocytes) and incubated for 30 min at room temperature. After three washes with antibody dilution buffer, cells were stained with either: 1) a FITC-labeled secondary antibody (CruzFluor™ 488, Santa Cruz Biotechnology, Cat. no. sc-516248) at 1:50 in antibody dilution buffer for primary cells, or 2) a PE-labeled secondary antibody (Goat anti-Mouse IgG (H+L), ThermoFisher, Cat. no. M30004-1) at 1:100 in antibody dilution buffer for lentivirus transduced EF1α-driven *CD46*-expressing cells (to avoid spectral overlap with eGFP). Both incubations were 30 min in the dark. Cells were then washed twice with PBS before flow cytometric analysis. Uninfected control cells were included in each experiment.

**Immunofluorescence Microscopy**

Localization of CD46 was determined by microscopy as previously described (2). In brief, cells were fixed with 4% PFA in PBS and blocked with 2% BSA in PBS. Cells were stained with a custom primary anti-bovine CD46 antibody followed by a goat anti-rabbit IgG FITC-conjugated secondary antibody (Abcam, catalog no. ab6717). Nuclei were counterstained with DAPI, and cells were visualized using an EVOS FL Auto microscope at 10x magnification (Thermo Fisher Scientific).

#### Monocyte and Lymphocyte Isolation and *Ex Vivo* BVDV Challenge

Peripheral blood mononuclear cells (PBMCs) were isolated from the edited and control animals and challenged with BVDV to assess virus resistance in a biologically relevant context. Blood was collected from the homozygous *CD46*-edited Gir heifer (Ginger), her heterozygous *CD46* offspring (Giraldo), and an age- and sex-matched control from a cross-bred beef calf (Bueno) via jugular venipuncture into syringes containing EDTA as an anticoagulant. Peripheral blood mononuclear cells (PBMCs) were isolated by density gradient centrifugation using SepMate tubes (Stemcell Technologies, Cambridge, MA, USA) (2). Monocytes were separated from lymphocytes by adherence selection, as previously described (2). Lymphocytes ($2.5x10^5$) were infected with BVDV as previously described with a maximal MOI (average MOI = 5) for each isolate based on well volume (6). Infection efficiency was quantified at 32 hours post-infection for lymphocytes by flow cytometric detection of the BVDV E2 glycoprotein (Creative Diagnostics; Catalog no. DMAB28412) (2, 6). For monocyte susceptibility assays, $4x10^5$ cells were challenged with BVDV isolates at an MOI of 0.01 or with a 1:10 dilution of serum collected from persistently infected cattle. Viral replication was quantified at 48 hpi for serum challenges and 72 hpi for isolate challenges by BVDV-specific RT-qPCR (7). The fold increase in viral RNA compared to the input concentration was determined with the delta Ct method and graphed with commercially available software (Prism, GraphPad Software, LLC, Boston, MA, USA).

**Long-Read Transcript Analysis of *CD46* Allele Expression and Isoform Selection**

Long-read transcript sequencing was used to evaluate *CD46* allele expression in cells derived from Giraldo and to identify the predominant *CD46* transcript isoform for use in the lentivirus complementation model. For Giraldo-derived samples, full-length multiplexed mRNA libraries were prepared from RNA extracted from whole blood and primary skin fibroblasts using the PacBio Iso-Seq/Kinnex workflow (Pacific Biosciences, Menlo Park, CA, USA) according to the manufacturer's protocol. Libraries were pooled and sequenced using proprietary reagents (Revio Polymerase Kit; Pacific Biosciences) across three Revio SMRT Cells with 24-hour data acquisition times.

To estimate the relative representation of edited and unedited *CD46* alleles in Giraldo's primary cells, full-length non-concatemer (FLNC) reads were screened for allele-specific sequence variation. Custom Perl scripts were developed to search sequencing files for a 19-nucleotide substitution corresponding to either the wild-type *CD46* allele (encoding $G_{82}$QVLAL) or the edited *CD46* allele (encoding $A_{82}$LPTFS). The scripts used Linux command-line utilities (gunzip and grep) to process a single skin fibroblast FASTQ file (12.5 GB) and two merged whole-blood FASTQ files (9.5 GB). The number of FLNC reads containing each allele-specific sequence was counted to estimate relative allele representation among *CD46* transcripts.

A separate analysis was performed to identify the predominant *CD46* transcript isoform expressed in MDBK cells for use in *CD46* complementation constructs. Previously generated PacBio Iso-Seq data from MDBK cells were reanalyzed for this purpose. Total RNA from MDBK cells was sequenced using the PacBio Iso-Seq platform in April 2021, and raw sequencing data were processed by the USMARC Core Lab using the PacBio Iso-Seq analysis pipeline to generate collapsed consensus transcript isoforms and associated full-length read counts. Consensus transcript sequences were imported into Geneious Prime and aligned to the longest annotated bovine *CD46* reference transcript (RefSeq accession no. NM_001242561) to identify the corresponding *CD46* transcript cluster. Isoform abundance was then determined based on the number of supporting full-length reads (count_fl) reported by the Iso-Seq pipeline.

#### ***TRIM5*** **Knockdown in MDBK-CD46Δ Cells**

An MDBK-based cell model was developed to evaluate the impact of individual *CD46* alleles on BVDV susceptibility. In this model, lentivirus transduction was used to express the *CD46* variants in a cell line lacking the entire *CD46* gene sequence (MDBK-CD46Δ; (2)). To facilitate efficient lentivirus integration, we used CRISPR/Cas9-mediated non-homologous end joining (NHEJ) to generate a *TRIM5* knockdown in the MDBK-CD46Δ cell line, as the *TRIM5* gene is a known retroviral restriction factor in bovine cells (8).

The gRNA sequence 5'-TTCTGTCAAGCCTGTATCAC-3' was designed using CHOPCHOP software v3 (9) to target exon two of the *TRIM5-3* (LOC505265) gene.  This specific paralog was targeted based on evidence that the *TRIM5* locus has undergone multiple rounds of duplication in cattle (10), whereas only the *TRIM5-3* gene has been shown to restrict retroviral infection in bovine cells (8).

Actively growing MDBK-CD46Δ cells were electroporated using the Neon NxT Electroporation System (Life Technologies; Carlsbad, CA, USA). For each electroporation, 100,000 cells were resuspended in Buffer GE and electroporated with 240 ng of gRNA and 1.25 μg of TrueCut Cas9 protein v2 (Invitrogen, Thermo Fisher Scientific, Waltham, MA, USA) using a 10 μL kit. Cells were electroporated using a single pulse at 1,200 V with a 40 ms pulse width and cultured in 500 μL of complete medium without antibiotics in a single well of a 24-well plate for 24 hours at 37°C in 5% $CO_2$. Media was replaced with 500 μL of complete medium after 24 hours.

After 48 hours, cells were sorted into single-cell clones using a Sony SH800 benchtop flow cytometer (Sony Biotechnology, San Jose, CA, USA). Individual cells were deposited into separate wells of a 96-well plate containing 100 μL of complete medium and incubated for 13 days to allow clonal expansion. Single-cell clones were then expanded to 24-well plates and incubated for 4 days.

**Genotyping *TRIM5*-edited Cells**

Cells from the 24-well plate were trypsinized and collected by centrifugation at 300 x g for 5 min at room temperature and the supernatant was removed. A fresh lysis buffer was prepared just before use by mixing a commercial cell lysis buffer (Invitrogen, Cat# 100034303) with Proteinase K in a ratio of 50 μL of buffer to 2 μL of Proteinase K per sample. The cell pellet was then resuspended in 52 μL of the prepared lysis buffer. The mixture was transferred to a PCR tube and incubated at 68°C for 15 min to promote cell lysis and protein degradation, followed by a 10 min incubation at 95°C to inactivate the Proteinase K. The DNA extract was then held at 4°C and either used immediately for PCR or stored at -20°C until use.

Putative knockout clones were screened by Illumina sequencing of the target amplicon, with libraries prepared using a two-step PCR protocol. The first PCR amplified the *TRIM5* locus using gene-specific primers that included Illumina bridge sequences (underlined): Forward, 5’-AGACGTGTGCTCTTCCGATCTACAAAGCCAACAAATGACCTTC-3’ and Reverse, 5’-TACACGACGCTCTTCCGATCTAGACAATAAGGAGCAGGTGAGC-3’. The PCR was performed with AmpliTaq Gold polymerase master mix (Thermo Fisher Scientific) under the following cycling conditions: an initial denaturation at 95°C for 10 min; 40 cycles of 95°C for 30 sec, 62.5°C for 30 sec, and 72°C for 30 sec; and a final extension at 72°C for 7 min. The PCR product was then purified by exonuclease treatment and ethanol precipitation.

The second PCR was performed to add unique indices to each sample. The reaction used a unique Illumina forward primer (5’-CAAGCAGAAGACGGCATACGAGAT*NNNNNN*GTGACTGGAGTTCAGACGTGTGCTCTTCCGATCT-3’) and a common reverse primer (5’-AATGATACGGCGACCACCGAGATCTACACTCTTTCCCTACACGACGCTCTTCCGATCT-3’). The PCR cycling conditions were as follows: an initial denaturation at 95°C for 10 min; 40 cycles of 95°C for 30 sec, 60°C for 30 sec, and 72°C for 30 sec; and a final extension at 72°C for 7 min. Amplicons were purified using a 0.7x concentration of AMPure XP beads (Beckman Coulter, Brea, CA, USA) and visualized by agarose gel electrophoresis. Libraries were quantified using a Qubit fluorometer, diluted to 4 nM, and sequenced on a MiSeq instrument using a 500-cycle MiSeq Reagent Nano v2 kit (Illumina, Inc.) with 2×250 base paired-end reads.

Amplicon sequencing data was analyzed using Geneious Prime (v.2025.0.2; Dotmatics, Boston, MA, USA). Adapter sequences and reads with Phred scores below 20 were trimmed using BBDuk (v.38.84). Short reads (<20 bp) were discarded, and the remaining paired reads were merged using BBmerge (11). The "Analyze CRISPR Editing Results" tool was used to identify modifications in the *TRIM5* sequence by comparing it to the reference amplicon sequence and the MDBK WGS.

Protein expression of TRIM5 could not be measured directly due to the lack of a commercial antibody that recognizes bovine TRIM5. Therefore, the knockdown was functionally confirmed by measuring the increase in lentivirus transgene expression (eGFP), which was necessary to achieve efficient transduction for our downstream experiments.

**Lentivirus Production**

To optimize transgene expression in target cells, promoter screening was initially conducted in MDBK-CD46Δ-TRIM5KD cells using the following transfer plasmids: pLJM1-EGFP (Cytomegalovirus [CMV] promoter; Addgene, Watertown, MA, USA, plasmid #19319), pTwist Lenti EF1α EGFP Puro (Elongation Factor 1-alpha [EF1α] promoter; Twist Bioscience, South San Francisco, CA, USA), pTwist Lenti SFFV Puro WPRE (Spleen Focus-Forming Virus [SFFV] promoter; Twist Bioscience), and pTwist Lenti CAG Puro (Chicken β-actin [CAG] promoter; Twist Bioscience). Based on transgene expression efficiency in MDBK-CD46Δ-TRIM5KD cells, the EF1α and SFFV promoters were selected for downstream applications expressing CD46 variants. Consequently, experimental transfer vectors consisted of either pWPI (serving as an eGFP control for screening cells for *TRIM5* knockout), or pTwist Lenti EF1α

EGFP Puro and pTwist Lenti SFFV Puro WPRE engineered to contain the *CD46* variant of interest.

For comparisons among MDBK *CD46*, the wild-type paternal Zacarias *CD46* allele, and the edited Ginger *CD46* allele, all rescue constructs were generated using the same *CD46* transcript isoform to ensure that differences in BVDV susceptibility reflected allelic variation rather than differences in *CD46* isoform structure. All constructs were synthesized and sequence-verified by Twist Bioscience.

For lentivirus production, HEK293T cells were seeded in 10 cm plates at a density calculated to achieve 70–80% confluence the following day. Prior to transfection, the cell culture medium was replaced with 9 mL of fresh DMEM supplemented with 10% FBS and 1× GlutaMAX, omitting antibiotic-antimycotic reagents. Transfections were performed using Lipofectamine 3000 Reagent (Thermo Fisher Scientific) according to the manufacturer's instructions. For each 10 cm plate, a third-generation packaging system was utilized consisting of a total of 20 μg of plasmid DNA: 4 μg pMD2.G, 3 μg pRSV-REV, 3 μg pMDLg/pRRE, and 10 μg of the respective transfer vector. The DNA-lipid complexes (1 mL) were added dropwise to the cells, and plates were incubated overnight at 37°C with 5% CO2.

Approximately 16 hours post-transfection, the transfection medium was removed and gently replaced with 10 mL of DMEM supplemented with 10% FBS, 1× GlutaMAX, and 5 mM sodium butyrate. Following an 8 hour incubation at 37°C with 5% CO2, the medium was exchanged for 5.5 mL of DMEM supplemented with 10% FBS, 1× GlutaMAX, and 10 mM HEPES. Cells were then incubated for an additional 24 hours under standard conditions to allow for viral production. Lentiviral supernatant (~5 mL per plate) was harvested approximately 24 hours after the HEPES media change, transferred to 15 mL conical tubes on ice, and clarified through a 0.45 µm syringe filter. The filtered lentivirus (~4.5 mL) was subsequently transferred to 100 kDa Amicon Ultra-15 centrifugal filter units (Millipore Sigma, St. Louis, MO, USA) and concentrated by centrifugation at 3,200 × *g* for 10 min at room temperature. The concentrated lentivirus was carefully collected and supplemented with polybrene at a ratio of 1 µL per 1 mL of concentrated virus prior to downstream application or storage.

**Lentivirus Transduction and Puromycin Selection**

MDBK-CD46Δ-TRIM5KD cells were seeded in 24-well plates to achieve 40% confluence the following day. The next day, the medium was removed and cells were inoculated with 500 µL of concentrated lentivirus. After 24 hours, the viral supernatant was removed, and the cells were incubated for 24 hours with fresh complete medium at 37°C in 5% $CO_2$. Two days after lentivirus transduction, puromycin selection was initiated. Cells were harvested using TrypLE Express (Gibco) and transferred from the 24-well plate to a 10 cm plate containing 9 mL of complete medium supplemented with 2 ug/mL puromycin. Transduced cells were maintained in 2 ug/mL puromycin until they were seeded for BVDV infection studies. CD46 expression and BVDV susceptibility were measured as described in the material and methods section "Flow Cytometric Quantification of CD46 Surface Expression and BVDV Infection."

### References


1. Heaton MP, Grosse WM, Kappes SM, Keele JW, Chitko-McKown CG, Cundiff LV, Braun A, Little DP, Laegreid WW. 2001. Estimation of DNA sequence diversity in bovine cytokine genes. Mamm Genome 12:32-7.
2. Workman AM, Heaton MP, Vander Ley BL, Webster DA, Sherry L, Bostrom JR, Larson S, Kalbfleisch TS, Harhay GP, Jobman EE, Carlson DF, Sonstegard TS. 2023. First gene-edited calf with reduced susceptibility to a major viral pathogen. PNAS Nexus 2:pgad125.
3. Heaton MP, Smith TP, Carnahan JK, Basnayake V, Qiu J, Simpson B, Kalbfleisch TS. 2016. Using diverse U.S. beef cattle genomes to identify missense mutations in EPAS1, a gene associated with pulmonary hypertension. F1000Res 5:2003.
4. Chakraborty R, Meagher TR, Smouse PE. 1988. Parentage analysis with genetic markers in natural populations. I. The expected proportion of offspring with unambiguous paternity. Genetics 118:527-536.
5. Workman AM, Heaton MP, Webster DA, Harhay GP, Kalbfleisch TS, Smith TPL, Falkenberg SM, Carlson DF, Sonstegard TS. 2021. Evaluating large spontaneous deletions in a bovine cell line selected for bovine viral diarrhea virus resistance. Viruses 13:2147.
6. Krueger AC, Vander Ley BL, Heaton MP, Sonstegard TS, Workman AM. 2025. Primary cells from a CD46-edited bovine heifer have reduced BVDV susceptibility despite viral adaptation to heparan sulfate. Viruses 17:634.

7. Mahlum CE, Haugerud S, Shivers JL, Rossow KD, Goyal SM, Collins JE, Faaberg KS. 2002. Detection of bovine viral diarrhea virus by TaqMan reverse transcription polymerase chain reaction. J Vet Diagn Invest 14:120-5.

8. Si Z, Vandegraaff N, O'Huigin C, Song B, Yuan W, Xu C, Perron M, Li X, Marasco WA, Engelman A, Dean M, Sodroski J. 2006. Evolution of a cytoplasmic tripartite motif (TRIM) protein in cows that restricts retroviral infection. Proc Natl Acad Sci U S A 103:7454-9.

9. Labun K, Montague TG, Krause M, Torres Cleuren YN, Tjeldnes H, Valen E. 2019. CHOPCHOP v3: expanding the CRISPR web toolbox beyond genome editing. Nucleic Acids Res 47:W171-W174.

10. Sawyer SL, Emerman M, Malik HS. 2007. Discordant evolution of the adjacent antiretroviral genes TRIM22 and TRIM5 in mammals. PLoS Pathog 3:e197.

11. Bushnell B, Rood J, Singer E. 2017. BBMerge - Accurate paired shotgun read merging via overlap. PLoS One 12:e0185056.

**Table S1. Bovine SNP List for 115 Highly Informative Parentage SNPs**

| Count | ARS-UCD1.2 Position | Variant alleles | Variant name | Variant alias |
|---|---|---|---|---|
| 1 | chr 1:3958867 | T/G | rs29012842 | ARS-USMARC-Parent-DQ381153 |
| 2 | chr 1:29963400 | A/G | rs29010795 | ARS-USMARC-Parent-DQ451555 |
| 3 | chr 1:58965351 | G/T | rs29012530 | ARS-USMARC-Parent-DQ404150 |
| 4 | chr 1:98519700 | T/C | rs42939801 | ARS-USMARC-Parent-DQ404149 |
| 5 | chr 1:126428831 | T/A | rs29003723 | ARS-USMARC-Parent-AY761135 |
| 6 | chr 1:149646917 | T/C | rs29019282 | ARS-USMARC-Parent-DQ404151 |
| 7 | chr 1:156749195 | A/G | rs17871566 | ARS-USMARC-Parent-AY943841 |
| 8 | chr 2:5374531 | C/T | rs29022245 | ARS-USMARC-Parent-DQ404152 |
| 9 | chr 2:26949128 | G/A | rs41257512 | ARS-USMARC-Parent-AY776154 |
| 10 | chr 2:45730559 | T/G | rs29003466 | ARS-USMARC-Parent-AY841151 |
| 11 | chr 2:64731387 | G/A | rs29011466 | ARS-USMARC-Parent-DQ422949 |
| 12 | chr 2:110396989 | T/C | rs29019900 | ARS-USMARC-Parent-DQ786757 |
| 13 | chr 3:3068311 | T/G | rs29012306 | ARS-USMARC-Parent-DQ422950 |
| 14 | chr 3:21144762 | C/T | rs29010510 | ARS-USMARC-Parent-DQ647187 |
| 15 | chr 3:40255504 | G/C | rs29001941 | ARS-USMARC-Parent-AY842472 |
| 16 | chr 3:49548031 | T/C | rs29001956 | ARS-USMARC-Parent-AY842473 |
| 17 | chr 3:51817697 | C/G | rs29003226 | ARS-USMARC-Parent-AY842474 |
| 18 | chr 3:57896539 | G/T | rs29010802 | ARS-USMARC-Parent-DQ435443 |
| 19 | chr 3:97595840 | C/T | rs29026932 | ARS-USMARC-Parent-DQ489377 |
| 20 | chr 3:115822760 | G/T | rs29012691 | ARS-USMARC-Parent-DQ839235 |
| 21 | chr 4:16215151 | C/T | rs29014143 | ARS-USMARC-Parent-DQ647186 |
| 22 | chr 4:20652215 | A/G | rs29002127 | ARS-USMARC-Parent-AY842475 |
| 23 | chr 4:93361477 | G/A | rs41590225 | ARS-USMARC-Parent-DQ485413 |
| 24 | chr 4:117385703 | G/A | rs29011099 | ARS-USMARC-Parent-DQ647188 |
| 25 | chr 5:7656578 | A/T | rs43710106 | ARS-USMARC-Parent-DQ470475 |
| 26 | chr 5:27670995 | A/G | rs43708500 | ARS-USMARC-Parent-DQ500958 |
| 27 | chr 5:62933839 | G/A | rs29012226 | ARS-USMARC-Parent-DQ647189 |
| 28 | chr 5:97631203 | G/A | rs17871338 | ARS-USMARC-Parent-AY844963 |
| 29 | chr 5:112616318 | T/G | rs17871378 | ARS-USMARC-Parent-DQ468384 |
| 30 | chr 5:118152314 | G/A | rs29023691 | ARS-USMARC-Parent-DQ846688 |
| 31 | chr 6:12732175 | C/T | rs29013632 | ARS-USMARC-Parent-DQ647190 |
| 32 | chr 6:45364629 | A/G | rs29017713 | ARS-USMARC-Parent-DQ789028 |
| 33 | chr 6:88813590 | A/G | rs41255758 | ARS-USMARC-Parent-AY849380 |
| 34 | chr 7:7248738 | C/A | rs29013741 | ARS-USMARC-Parent-DQ888309 |
| 35 | chr 7:17223080 | C/A | rs29024430 | ARS-USMARC-Parent-DQ786758 |
| 36 | chr 7:53233377 | G/A | rs29012174 | ARS-USMARC-Parent-DQ650635 |
| 37 | chr 7:79304309 | A/T | rs29009979 | ARS-USMARC-Parent-DQ916057 |

| | | | | |
|---|---|---|---|---|
| 38 | chr 7:91858328 | T/C | rs29026696 | ARS-USMARC-Parent-DQ786759 |
| 39 | chr 8:1688535 | A/G | rs3423092698 | ARS-USMARC-Parent-DQ916058 |
| 40 | chr 8:28747847 | A/G | rs29024525 | ARS-USMARC-Parent-DQ650636 |
| 41 | chr 8:59585336 | C/T | rs41255750 | ARS-USMARC-Parent-AY850194 |
| 42 | chr 8:87523043 | G/T | rs29010468 | ARS-USMARC-Parent-DQ837644 |
| 43 | chr 8:104418638 | T/C | rs29011266 | ARS-USMARC-Parent-DQ674265 |
| 44 | chr 9:45135255 | A/G | rs29011985 | ARS-USMARC-Parent-DQ846689 |
| 45 | chr 9:97029776 | A/C | rs29009858 | ARS-USMARC-Parent-DQ786765 |
| 46 | chr 10:3582438 | G/A | rs29012070 | ARS-USMARC-Parent-DQ786766 |
| 47 | chr 10:14700394 | T/G | rs3423302605 | ARS-USMARC-Parent-DQ786760 |
| 48 | chr 10:44059494 | C/T | rs29012840 | ARS-USMARC-Parent-DQ786761 |
| 49 | chr 10:55539558 | A/G | rs29012019 | ARS-USMARC-Parent-DQ984827 |
| 50 | chr 10:81223782 | G/T | rs29010772 | ARS-USMARC-Parent-DQ786762 |
| 51 | chr 10:97152695 | G/A | rs29012457 | ARS-USMARC-Parent-DQ984825 |
| 52 | chr 11:1729993 | C/G/T | rs29012894 | ARS-USMARC-Parent-DQ837646 |
| 53 | chr 11:24487930 | T/C | rs29015870 | ARS-USMARC-Parent-DQ837645 |
| 54 | chr 11:46544231 | T/C | rs41255717 | ARS-USMARC-Parent-AY851162 |
| 55 | chr 11:66365240 | A/G | rs29018818 | ARS-USMARC-Parent-DQ837643 |
| 56 | chr 11:103007477 | T/C | rs17871661 | ARS-USMARC-Parent-AY851163 |
| 57 | chr 12:11793754 | T/A | rs29020472 | ARS-USMARC-Parent-DQ786763 |
| 58 | chr 12:76649683 | C/A | rs29012872 | ARS-USMARC-Parent-DQ832700 |
| 59 | chr 13:2079650 | T/C | rs29011643 | ARS-USMARC-Parent-EF026087 |
| 60 | chr 13:25335505 | A/T | rs29009668 | ARS-USMARC-Parent-EF034081 |
| 61 | chr 13:47037532 | G/A | rs41257524 | ARS-USMARC-Parent-AY853302 |
| 62 | chr 13:74664505 | A/G | rs41257490 | ARS-USMARC-Parent-AY853303 |
| 63 | chr 14:9116686 | A/G | rs43710049 | ARS-USMARC-Parent-DQ846690 |
| 64 | chr 14:26068764 | T/A | rs29027559 | ARS-USMARC-Parent-DQ984826 |
| 65 | chr 14:46226278 | C/T | rs29019814 | ARS-USMARC-Parent-DQ846691 |
| 66 | chr 14:77723052 | T/C | rs29010281 | ARS-USMARC-Parent-DQ846692 |
| 67 | chr 15:20899423 | C/T | rs43708441 | ARS-USMARC-Parent-EF042090 |
| 68 | chr 15:37547916 | G/A | rs43706884 | ARS-USMARC-Parent-DQ866817 |
| 69 | chr 15:77950839 | C/A | rs29011701 | ARS-USMARC-Parent-DQ866818 |
| 70 | chr 16:9259631 | T/C | rs29017621 | ARS-USMARC-Parent-DQ846693 |
| 71 | chr 16:32760431 | A/C | rs29015842 | ARS-USMARC-Parent-DQ846694 |
| 72 | chr 16:66121452 | G/C/T | rs17871214 | ARS-USMARC-Parent-AY857620 |
| 73 | chr 16:78825060 | T/C | rs41825023 | ARS-USMARC-Parent-DQ846695 |
| 74 | chr 17:1023299 | T/G | rs29012422 | ARS-USMARC-Parent-DQ888310 |
| 75 | chr 17:29524233 | C/G | rs29002256 | ARS-USMARC-Parent-AY858890 |
| 76 | chr 18:1801945 | C/T | rs29014953 | ARS-USMARC-Parent-EF028073 |
| 77 | chr 18:23351423 | C/T | rs29009907 | ARS-USMARC-Parent-DQ916059 |

| | | | | |
|---|---|---|---|---|
| 78 | chr 18:48545897 | C/A | rs17871403 | ARS-USMARC-Parent-AY914316 |
| 79 | chr 19:8274910 | G/C/T | rs29017313 | ARS-USMARC-Parent-DQ888311 |
| 80 | chr 19:15017885 | A/G | rs29025380 | ARS-USMARC-Parent-EF026084 |
| 81 | chr 19:35808547 | G/A | rs29015945 | ARS-USMARC-Parent-DQ888312 |
| 82 | chr 19:44173520 | C/T | rs41257460 | ARS-USMARC-Parent-AY916666 |
| 83 | chr 19:54554240 | T/C | rs29011141 | ARS-USMARC-Parent-EF164803 |
| 84 | chr 20:788373 | T/C | rs29010004 | ARS-USMARC-Parent-DQ984828 |
| 85 | chr 20:17847390 | A/G | rs43708490 | ARS-USMARC-Parent-DQ888313 |
| 86 | chr 20:36553740 | C/G | rs29012811 | ARS-USMARC-Parent-DQ990835 |
| 87 | chr 21:3040671 | A/G | rs43706859 | ARS-USMARC-Parent-DQ995976 |
| 88 | chr 21:26120612 | G/T | rs29012316 | ARS-USMARC-Parent-EF093511 |
| 89 | chr 21:63550319 | G/T | rs29021607 | ARS-USMARC-Parent-EF026085 |
| 90 | chr 22:11000418 | G/A | rs29015065 | ARS-USMARC-Parent-DQ990832 |
| 91 | chr 22:22468414 | T/A | rs29015170 | ARS-USMARC-Parent-EF093509 |
| 92 | chr 22:45656061 | A/C | rs29010035 | ARS-USMARC-Parent-EF093510 |
| 93 | chr 22:55883157 | C/T | rs29013532 | ARS-USMARC-Parent-EF034082 |
| 94 | chr 23:7318640 | A/G | rs41255852 | ARS-USMARC-Parent-AY929334 |
| 95 | chr 23:27495679 | A/G | rs17872223 | ARS-USMARC-Parent-AY937242 |
| 96 | chr 23:51029219 | T/A | rs29020870 | ARS-USMARC-Parent-EF089234 |
| 97 | chr 24:1617199 | G/A | rs3423094146 | ARS-USMARC-Parent-DQ995977 |
| 98 | chr 24:15141986 | A/G | rs29010147 | ARS-USMARC-Parent-DQ990833 |
| 99 | chr 24:55950028 | T/C | rs17870274 | ARS-USMARC-Parent-AY939849 |
| 100 | chr 25:3130565 | C/A | rs29018286 | ARS-USMARC-Parent-EF034083 |
| 101 | chr 25:14593246 | T/C | rs17872131 | ARS-USMARC-Parent-AY941204 |
| 102 | chr 25:40306557 | A/G | rs29003010 | ARS-USMARC-Parent-AY942198 |
| 103 | chr 26:8192400 | T/C | rs29013727 | ARS-USMARC-Parent-DQ990834 |
| 104 | chr 26:13194710 | A/G | rs29023666 | ARS-USMARC-Parent-EF150946 |
| 105 | chr 26:37900334 | C/T | rs43708440 | ARS-USMARC-Parent-EF034086 |
| 106 | chr 27:16084911 | C/G | rs29013546 | ARS-USMARC-Parent-EF093512 |
| 107 | chr 27:22410377 | C/G | rs29016185 | ARS-USMARC-Parent-EF034084 |
| 108 | chr 27:37824983 | T/C | rs29015783 | ARS-USMARC-Parent-EF141102 |
| 109 | chr 28:5874287 | A/G | rs29025677 | ARS-USMARC-Parent-EF034085 |
| 110 | chr 28:15972475 | G/A | rs29017064 | ARS-USMARC-Parent-EF034087 |
| 111 | chr 28:35132199 | T/C | rs29013660 | ARS-USMARC-Parent-EF026086 |
| 112 | chr 28:43879624 | C/T | rs29014974 | ARS-USMARC-Parent-EF042091 |
| 113 | chr 29:9104654 | A/G | rs17871190 | ARS-USMARC-Parent-AY856094 |
| 114 | chr 29:28278699 | C/T | rs29024749 | ARS-USMARC-Parent-EF034080 |
| 115 | chr 29:44090714 | A/G | rs110843280 | ARS-USMARC-Parent-DQ404153 |

**File S2. *CD46* Open Reading Frame Sequences used for Lentivirus Complementation Studies**

### MDBK CD46 ORF

ATGAGGGCGTCTTGCACCCCGCTGAAGGCGCCGCTCCGCCGCCCCGAAAGACTGGCTTCTTCTGGGCGCTTCGCCTGGGT
GCTTCTGCTGGCGCCGCTGCTCCTGCTGCCCACGTCCTCCGATGCCTGTGATGATCCACCAAGATTTGTCTCTATGAAGC
CCCAGGGTACCCTTAAACCCAGTTATAGTCCTGGGGAGCAGATTGTGTATGAATGTCGTCTGGGTTTCCAGCCAATAACT
CCTGGTCAAGTCCTGGCTCTCGTTTGTCAGGATAATAATACATGGTCGTCTCTCCAGGAGGGCTGTAAAAAAAGACGGTG
TCCTACCCTAGCTGATCCCACAAATGGCCAAGTTATCCTTGTAAATGGAAGTACTGAGTTTGGCTCAGAGGTTCACTATG
TTTGTAATAATGGTTATTACTTACTGGGGACAAATATTTCTTATTGTGAAGTTTCTTCTGGAACTGGTGTGAACTGGAGT
GATAATCCTCCAACATGTGAAAAGATTTTGTGTCAACCGCCTCCAGAAATTCAAAATGGAAAATACACCAATAACCACAA
GGATGTATTTGAATACAATGAAGTAGTAGCTTATAGTTGTGATCCTTCAAATGGGCCAGATGAATATTCCCTTGTTGGAG
AGAGCAAGCTTACTTGTATTGGAAATGGTGAATGGAGTAGTCAACCCCCTCAGTGTAAAGTGGTCAAATGTGTATATCCA
GCCATTGAACATGGAACGATAGTCTCAGGATTTGGACCAAAATATTACTACAAAGCGACGGTTGTACTTAAATGCAATGA
GGGTTTTAACCTTTATGGCAACAGTGTAGTTGTCTGTGGTGAGAACAGTACTTGGGAGCCCGAGCTACCAAAGTGTATTA
AAGGACATCCCCCCCGTCCTACTGATGCATCACCCCCTAACGGTGCTGAGGGTTTAGGTGCAGGATACATCGTGCTCGTC
ATTGTTGCTGTACTTATTGGCGTTGGATTATTGCTCTGCCTGTACTGCTGTTTTTGCAGACAGAGGAAGAAAGGGAAAGC
AGAATGTAGCGCTACGTACACCACTTATCAGGATAAAGCAACCACTGCAACAGAACAGATGAACTGA

### Zacarias CD46 ORF

ATGAGGGCGTCTTGCACCCCGCTGAAGGCGCCGCTCCGCCGCCCCGAAAGACTGGCTTCTTCTGGGCGCTTCGCCTGGGT
GCTTCTGCTGGCGCCGCTGCTCCTGCTGCCCACGTCCTCCGATGCCTGTGATGATCCACCAAGATTTGTCTCTATGAAGC
CCCAGGGTACCCTTAAACCCAGTTATAGTCCTGGGGAGCAGATTCTGCATGAATGTCGTCCGGGGTTCCAGCCAATAACT
CCTGGTCAAGTCCTGGCTCTCGTTTGTCAGGATAATAATACATGGTCGTCTCTCCAGGAGGGCTGTAAAAGAAGACAGTG
TCCTAACCTAGCTGATCCCACAAATGGCCAAGTTATCCTTGTAAATGGAAATACTGAGTTTGGCTCAGAGGTTCACTATG
TTTGTAATAATGGTTATTACTTACTGGGGACAAATATTTCTTATTGTGAAGTTTCTTCTGGAACTGGTGTGAACTGGAGT
GATAATCCTCCAACATGTGAAAAGATTTTGTGTCAACCACCTCCAGAAATTCAAAATGGAAAATACACCAATAGCCACAA
GGATGTATTTGAATACAATGAAGTAGTAACTTATAGTTGTGATCCTTCAAATGGACCAGATGAATATTCACTTGTTGGAG
AGAGCAAGCTTACTTGTATTGGAAATGGTGAATGGAGTAGTAAACCCCCTCAGTGTAAAGTGGTCAAATGTGTATATCCA
GCCATTGAATATGGAACGATAGTCTCAGGATTTGGACCAAAATATTACTACAAAGCGACGGTTGTACTTAAATGCAATGA
GGGTTTTAACCTTCAAGGCAACAGCGTAGTTGTCTGTGGTGAGAACAGTACTTGGGAGCCCGAGCTACCAAAGTGTATTA
AAGGACATCCCCCCCGTCCTACTGATGCATCACCCCCTAACGGTGCTGAGGGTTTAGGTGCAGGATACATCGTGCTCGTC
ATTGTTGCTGTACTTATTGGCGTTGGATTATTGCTCTGCCTGTACTGCTGTTTTTGCGGACAGAGGAAGAAAGGGAAAGC
AGAATGTAGCGCTACGTACACCACTTATCAGGATAAAGCAACCACTGCAACAGAACAGATGAACTGA

### Ginger CD46 ORF

ATGAGGGCGTCTTGCACCCCGCTGAAGGCGCCGCTCCGCCGCCCCGAAAGACTGGCTTCTTCTGGGCGCTTCGCCTGGGT
GCTTCTGCTGGCGCCGCTGCTCCTGCTGCCCACGTCCTCCGATGCCTGTGATGATCCACCAAGATTTGTCTCTATGAAGC
CCCAGGGTACCCTTAAACCCAGTTATAGTCCTGGGGAGCAGATTGTGTATGAATGTCATCTGGGTTTCCAGCCAGTAACT
CCGGCATTGCCTACCTTCAGTGTTTGTCAGGATAATAATACATGGTCGTCTCTCCAGGAGGGCTGTAAAAAAAGACGGTG
TCCTACCCTAGCTGATCCCACAAATGGCCAAGTTATCCTTGTAAATGGAAATACTGAGTTTGGCTCAGAGGTTCACTATG
TTTGTAATAATGGTTATTACTTACTGGGGACAAGTATTTCTTATTGTGAAGTTTCTGGAAATGGTGTGAACTGGAGTGAT
AATCCTCCAACATGTGAAAAGATTTTGTGTCAACCGCCTCCAGAAATTCAAAATGGAAAATACACCAATAGCCACAAGGA
TGTATTTGAGTACAATGAAGTAGTAACTTATAGTTGTGATCCTTCAAATGGGCCAGATGAATATTCCCTTGTTGGAGAGA
GCAAGCTTACTTGTATTGGAAATGGTAAATGGAGTAGTCAACCCCCTCAGTGTAAAGTGGTCAAATGTGTATATCCAGCC
ATTGAATATGGAACGATAGTCTCAGGATTTGGACCAAAATATTACTACAAAGCGACGGTTGTACTTAAATGCAATGAGGG
TTTTAACCTTCAAGGCAACAGCGTAGTTGTCTGTGGTGAGAACAGTACTTGGGAGCCCGAGCTACCAAAGTGTATTAAAG
GACATCCCCCCCGTCCTACTGATGCATCACCCCCTAACGGTGCTGAGGGTTTAGGTGCAGGATACATCGTGCTCGTCATT
GTTGCTGTACTTATTGGCGTTGGATTATTGCTCTGCCTGTACTGCTGTTTTTGCGGACAGAGGAAGAAAGGGAAAGCAGA
ATGTAGCGCTACGTACACCACTTATCAGGATAAAGCAACCACTGCAACAGAACAGATGAACTGA
